# ARCHIVE: An Efficient Open-Ended DNA Recording Device Capable of Multiplex Capture of Pol-II Transcribed Signals

**DOI:** 10.64898/2026.07.10.737758

**Authors:** Aaron H. Rosenstein, Michael Garton

## Abstract

Engineering cell-based devices to record events into DNA has potential both as a non-ablative research tool and for enacting gene-circuit-based logic of cell therapies conditional on cell history. Whether as a means of understanding interactions on the single-cell level, or reconstructing histories of cellular events, a cellular DNA recording device has widespread utility, with prime editing-based methods at the forefront of this endeavor – notably peCHYRON. Open-ended recording tool resolution is inherently constrained by edit insertion efficiency however, and cannot yet capture RNA-polymerase II-transcribed signals, which constitute essentially all nuclear mammalian protein coding genes. To address this, we developed ARCHIVE (Amplified Recording of Cellular Histories into Information-dense Vectors of Events) by using machine-learning assisted prediction of prime-editing efficiency as a surrogate fitness model for generative *in silico* pegRNA evolution. ARCHIVE is a recording module capable of integrating RNA-encoded signals into predefined genomic loci with unprecedented efficiency and an order-of-magnitude improvement in temporal resolution.

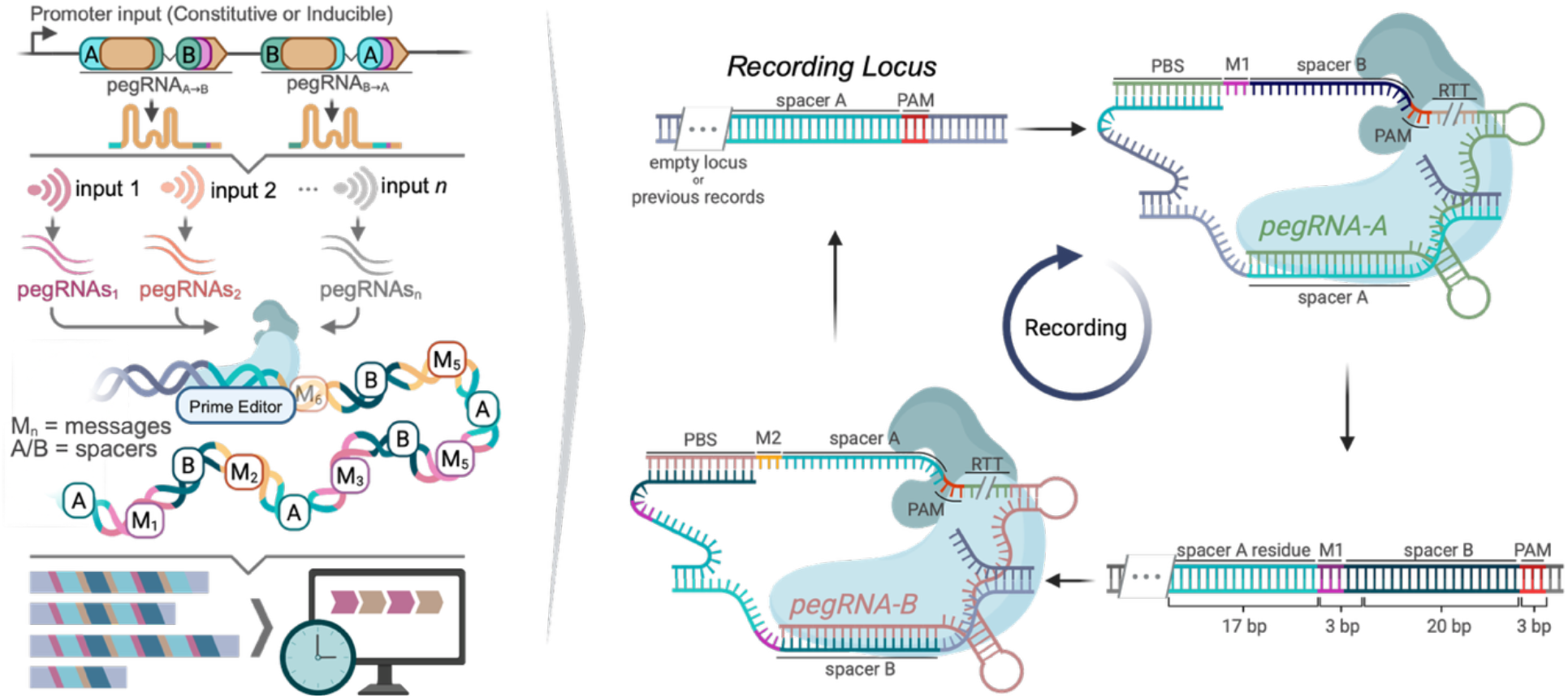

## Introduction

Engineering cells to act as miniature computer systems is an overarching goal of synthetic biology, paving the way for smarter precision cell therapeutics and diagnostics^1–3^. Many cellular devices have been developed which act as analogues to logic gates, amplifiers, oscillators, sensors, data recorders, and other well-characterized components of computer/robotic systems^4–8^. Cellular recording, a central concept to cellular computation, involves retention of information regarding cellular events for later retrieval by investigators or for conditional actuation by the cell. Such devices have utilized RNA, DNA and protein as recording substrates in both bacterial and mammalian cells^5,9,10^.

While both RNA and protein recorders are effective at capturing short-term event memory, their transient nature is suboptimal to DNA recorders, where memory can be retained in perpetuity. Other mechanisms of recording cell information such as single-cell RNA-sequencing (scRNA-Seq) are ablative, and thus cannot continuously record events^5^. Live-cell imaging can non-disruptively measure cellular phenomena continuously, but is not multiplexable beyond the number of fluorescent inputs, and is incompatible with *in-vivo* studies^11^. As such, recorders which integrate time-ordered orthogonal inputs into the genome are highly desirable, presenting a means of acquiring previously inaccessible information for potential research and cell-therapeutics use.

DNA recording systems were initially developed in bacteria. SCRIBE uses retron-encoded ssDNA templates to drive genomic integration of recorded information without temporal information ^12^. TRACE and Record-Seq use Cas1-Cas2 to chronologically integrate signals into an engineered CRISPR-array; it has not been shown that either function heterologously in mammalian systems ^13–15^. Most eukaryotic DNA recording devices rely on Cas9 as the central mediator, with BLADE being a noteworthy recombinase-based exception^16^. GESTALT, MEMOIR, mSCRIBE, and CHYRON use random mutagenesis to capture event magnitude^17–20^, while mCAMERA and INSCRIBE utilize base editing to modify predefined genomic loci^21,22^. INSCRIBE most recently demonstrated the capability to record signal magnitude with an in-situ readout. While alternate approaches to cell history recording include both protein-based oligomerization and intracellular RNA capture in vault proteins^23,24^, a structural limitation of previous DNA based systems is that none are yet capable of recording multiple events in chronological order.

Two recording systems, the ‘DNA Typewriter’ and peCHYRON have leveraged the sequence-specific DNA-writing capabilities of prime editing to record signals into defined recording loci^25,26^. Both systems present major advances in non-ablative cell measurement technology. DNA typewriter has been implemented for both lineage tracing and event recording^26–29^ while peCHYRON has been used to record cis-regulatory element activity and cell-signaling events from RNA polymerase II (Pol-II) promoter architectures. Whereas ‘DNA Typewriter’ integrates a defined number of events into a ‘DNA tape’ of fixed length preinserted into the genome, peCHYRON can be propagated *ad infinitum,* representing the state of the art in open-ended recorder design.

In both systems, the efficiency of each successive recording step limits the attainable temporal resolution and requires optimisation. We hypothesised that optimisation of temporal resolution coupled with extension of recording to the Pol-II machinery would enable robust capture of native gene-expression signals.

Here we develop ARCHIVE (Amplified Recording of Cellular Histories into Information-dense Vectors of Events). Building on the peCHYRON platform, ARCHIVE is a prime-editing-based recording architecture capable of efficiently capturing gene expression signals into genomic DNA (Figure 1a,b). For each input, two pegRNAs are expressed which constitute a cyclical prime-editing targeting scheme. PegRNA_A◊B_ targets spacer A, catalyzing templated message integration followed by spacer B. PegRNA_B◊A_ targets spacer B, catalyzing templated message integration followed by chaining of an additional spacer A, enabling further message integration events (Figure 1b). Both pegRNAs are linked to an expression input, enabling open-ended recording of pegRNA sequences tied to cellular event occurrences.

**Figure 1:**
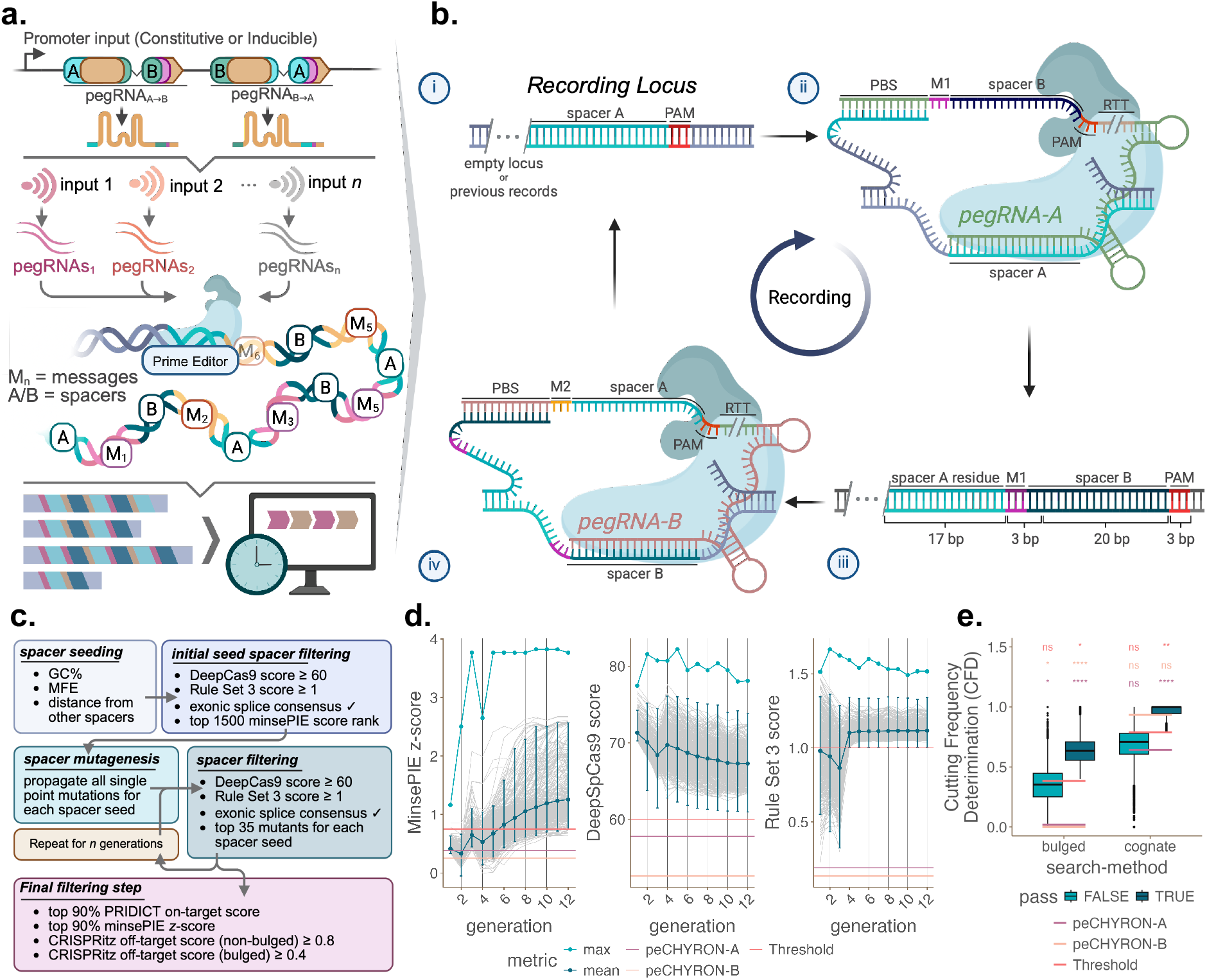
ARCHIVE Recording Paradigm, Mechanism, and *in Silico* Design. **(a)** Input of interest drives polycistronic expression of two intron-split pegRNAsto improve iterative efficiency. Distinct pegRNA-encoded trinucleotide message sequences delineate input identity, constituting an ordered record of events into a genomic recording locusfor subsequent retrieval by NGS. **(b)** i. Recording locus ground-state, where spacer A may be preceded by additional records. ii. Targeting by pegRNA_A◊B_ catalyzes integration of message 1 (M1) and spacer B. iii. Recording locus state after first editing-step. iv. Subsequent targeting of spacer B by pegRNA _B◊A_ inserts a second message (M2) followed by spacer A, reconstituting the original recording locus state for further targeting. **(c)** Genetic algorithm-based approach for spacer evolution with high Cas9-targeting and insertion efficiency by prime-editing. **(d)** MinsePIE, DeepSpCas9, and RS3 scoring over 12 generations of evolution. Teal points represent maximum spacer score, while navy represents mean score. Gray lines represent individual mean trajectories in score for each spacer cluster. Both peCHYRON spacer scores in addition to thresholds for inclusion in subsequent analyses are depicted. **(e)** CRISPR off-target scoring using both fully-cognate and single-bulge approaches using CRISPRitz. Significance of passing/failing guide distributions for both bulged and cognate off-target analyses was calculated with a one-sided asymptotic permutation test against the peCHYRON-A/B benchmarks and pass-threshold (significance text is coloured based on comparison), and adjusted with Benjamini-Hochberg (BH) correction. α=0.05. Statistical significance: **** p < 0.0001, *** p < 0.001, ** p < 0.01, * p < 0.05.

PegRNA design is critical for high-efficiency prime editing^30–33^ –– a challenge amplified in this case by the requirement for compatible sets of reciprocating pegRNAs (A ◊ B, B ◊ A). ARCHIVE leverages machine-learning-based models for prime-editing efficiency prediction from pegRNA sequence to develop reciprocating pegRNAs optimized for improved recording rates. Novel designs also facilitated avoidance of pegRNA expression cassette corruption by off-target prime-editing. These advances increase recording resolution, simplify pegRNA expression topology, and enable capture of RNA polymerase II-driven inputs, effectively increasing recording efficiency and enhancing potential for portability across input modalities.

## Results

### Machine-Guided Evolution of Spacers to permit simultaneous On-Target Prime Editor Targeting and Edit Integration

In silico design of pegRNAs was performed using a surrogate-assisted genetic algorithm to generate pairs of reciprocating pegRNAs with high iterative efficiencies (Figure 1c,d). Editing by pegRNA_A◊B_ licenses spacer-B insertion, which enables subsequent editing by pegRNA_B◊A_. MinsePIE and PRIDICT were employed as fitness predictors to direct evolution of 20-nt pegRNA spacer sequences with high prime editing efficiency for message+spacer insertion, whilst thresholding on Cas9 targeting efficiency (determined as Rule Set 3 (RS3) z-score ≥ 1 and DeepSpCas9 ≥ 60). All spacers were required to retain a CCT sequence, the reverse complement of the exonic splice consensus. Because the insert–homology-arm portion, not the PBS, carries MinsePIE’s predictive signal, edit-efficiency predictions varied homology-arm length at a fixed HEK3 target, and successive rounds of mutagenesis increased predicted insertion scores (Figure 1d). Top candidates (MinsePIE z-score ≥ 0.75) were scored for off-target potential using CRISPRitz (Figure 1e). Finally, an additional round of prime-editing efficiency prediction was conducted with PRIDICT, with top spacer candidates selected based on a combination of thresholds to sample a range of model-performance vs. genomic-specificity trade-offs (see Methods). Minimal correlation was observed between metrics (Supplementary Figure 1a).

### Experimental Validation and Optimization of Spacer Integration

Top-performing spacer sequences were concatenated with a degenerate 3-nt message sequence and ordered as an oligonucleotide library, cloned into a HEK3-targeting pegRNA destination vector and transfected into HEK293T cells to quantify spacer insertion rates (Figure 2a). RTT homology arm (HA) lengths from 26–50 nt (4-nt increments) were tested for each spacer. The pegRNA library was co-transfected with pPRG6-CMV-PE2-P2A-EGFP for three days, followed by FACS to isolate the EGFP+ cell fraction and targeted Illumina-sequencing of the HEK3 genomic locus. MinsePIE showed low-moderate in-sample predictive accuracy for HA lengths ≤ 38 nt (R^2^ = 0.21–0.51) and minimal correlation above this threshold (Supplementary Figure 1b). To improve prediction beyond the MinsePIE z-score baseline, a random forest model was trained to predict integration efficiency based on pegRNA sequence information using empirical editing results obtained from pegRNA Library 1. Performance was quantified as the R² between predicted and measured integration efficiency (Figure 2b). The random forest improved in-sample accuracy to R^2^ ≈ 0.7 from R^2^ ≈ 0.42 using MinsePIE alone, with generalization estimated by spacer-stratified 5-fold cross-validation (R2 ≈ 0.51) to prevent leakage between RTT-length replicates of the same spacer. Additional informative features were identified: MFE of the RTT, the nucleotide at spacer position 15, PRIDICT on-target efficiency, and HA length (Supplementary Figure 1c). Like MinsePIE, the random forest remained predictive only for shorter HA lengths (Supplementary Figure 1d). This informed the design of a second spacer library, which yielded spacers with further increases in insertion efficiency (Figure 2c,d).

**Figure 2:**
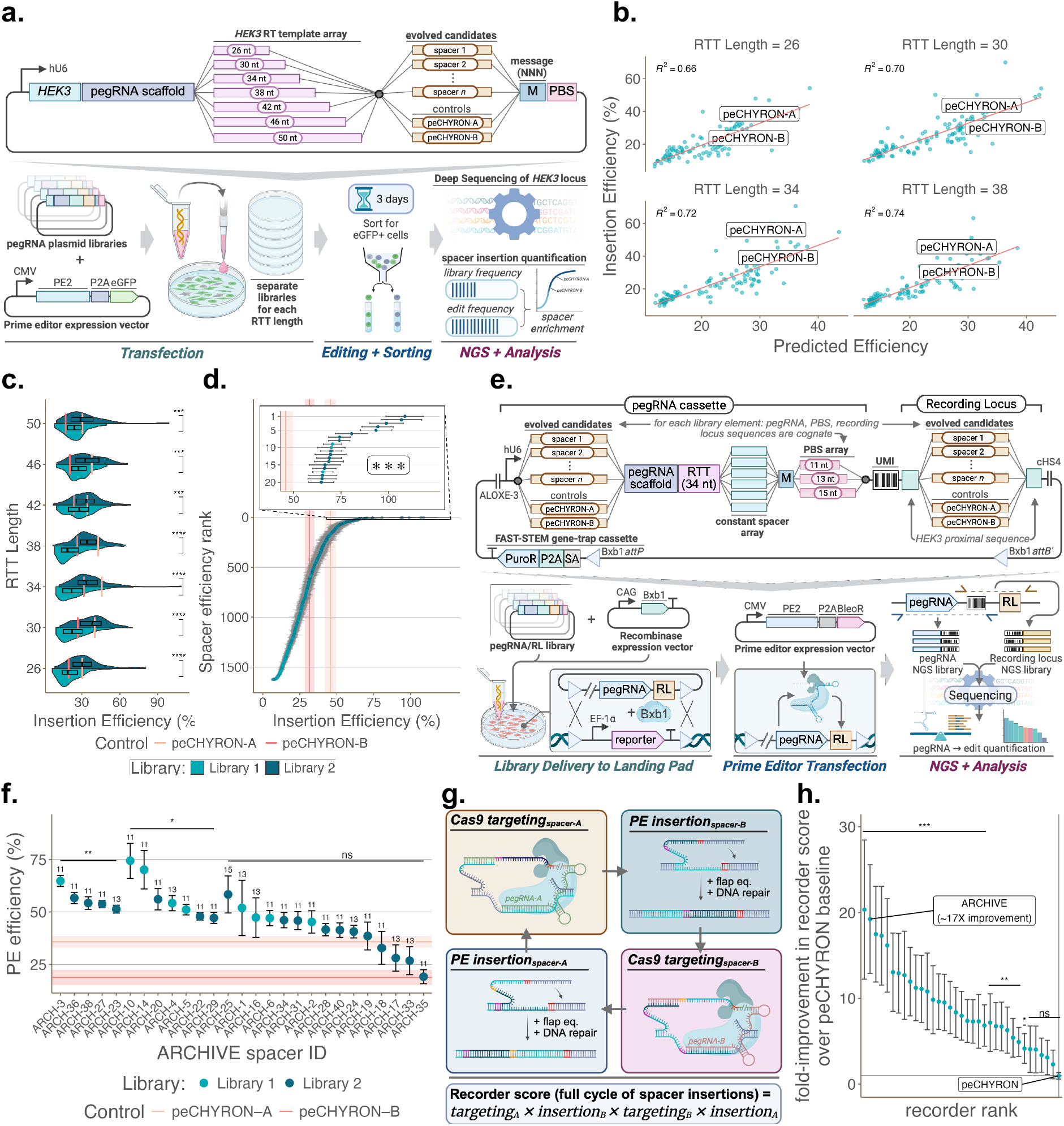
Experimental Validation of Generative pegRNA Design for Improved Recording Efficiency. **(a)** Pooled spacer insertion validation. pegRNA design depicts variable RTT lengths targeting *HEK3* for insertion of the spacer library. The pegRNA library was transfected with pPRG6-CMV-PE2-P2A-EGFP. Cells were sorted for EGFP+ status followed by deep-sequencing of the *HEK3* locus and pegRNA library for comparative spacer enrichment quantification to identify efficiently-inserted spacer library elements. **(b)** Random-forest model comparison between predicted and experimental spacer insertion efficiency for RTT lengths ≤ 38 nt. Linear regression model and R^2^ coefficient of determination are overlayed. Both peCHYRON spacers are overlayed (RTT length = 34 nt was used in original peCHYRON study). **(c)** Split-violin-plot with inline boxplot comparing spacer insertion efficiencies across both pegRNA libraries segmented by RTT-HA length. peCHYRON spacer efficiencies are overlayed. Library distributions for each RTT length were compared by two-sided Wilcoxon-Mann-Whitney tests followed by BH correction **(d**) Spacer ranking based on insertion efficiency. Error bars represent standard error of the mean across all trinucleotide message identities (nominal n = 50 trinucleotides / spacer). Mean (line) and standard error (shading) of peCHYRON-A/B control spacers are overlayed. Top 20 spacer efficiencies are depicted in inset plot. Efficiency for all generated spacers was compared to the peCHYRON-A control spacer using a bootstrap-permutation test (20,000 iterations). To account for the pooled screen design, resampling was weighted by the relative read-abundance of each trinucleotide message. p-values were adjusted using the BH correction. All comparisons in inset-plot were significant (***) **(e)** Pooled spacer targeting efficiency validation. PegRNA cassette map with cognate target site is depicted, cloned into a FAST-STEM-compatible delivery vector. Spacers and PBS elements are cognate for each library element. For each spacer, three PBS lengths (11,13,15 nt) were assessed to insert six possible spacer-length sequences. PegRNA library was stably integrated into HEK293T landing-pad cells by Bxb1-mediated recombination, followed by transfection pCMV-PE2-P2A-BleoR to induce editing. Both pegRNA and target site loci were amplified alongside UMI sequences for *in silico* association following Illumina-sequencing to quantify pegRNA identities driving efficient targeting to a recording locus. **(f)** Editing efficiency afforded by most-optimal PBS (labelled above points) for each candidate spacer. Error bars represent standard error of the mean across six insertion sequence identities. Mean (line) and standard error (shading) of peCHYRON-A/B control spacers are overlayed. Editing efficiencies for all spacers were compared to peCHYRON-A control with an unpaired one-sided t-test followed by BH correction. PBS lengths for each spacer which maximized insertion efficiency are reported above each error bar. **(g)** Recorder score calculation using a four-element product. targeting*_x_* = spacer *x* targeting efficiency. insertion*_x_* = spacer *x* insertion efficiency. **(h)** Comparison of top recorder scores to the peCHYRON control. Mean scores and standard errors (SE) were estimated by resampling (20,000 iterations) across four-element products. Statistical significance was determined by a weighted bootstrap hypothesis test comparing each candidate’s distribution to the peCHYRON control, with p-values adjusted via BH correction. α=0.05. Statistical significance: **** p < 0.0001, *** p < 0.001, ** p < 0.01, * p < 0.05.

### Experimental Validation of Spacer Targeting

On-target prime-editing efficiencies of spacers (each of which was independently efficient for HEK3 integration) were quantified by pooled screening in a HEK293T landing-pad cell line (HEK293T-LP; Figure 2e). Each library variant was constructed as a single DNA molecule encoding both a pegRNA cassette and its cognate target site. For every spacer, three PBS lengths (11, 13, 15 nt) and six spacer-length insertion sequences were tested combinatorially. Subsequent transfection of the prime editor and amplification of the pegRNA–target locus permitted simultaneous determination of pegRNA identity and editing efficiency, with 12 spacers exhibiting significant increases in integration efficiency compared to the peCHYRON-A control sequence (47%–74% vs. 35%; Figure 2f), while peCHYRON-B exhibited a nominal 18% integration efficiency.

To identify the two spacers that, when deployed in a reciprocating recording scheme, enabled the highest sequential recording efficiency, a theoretical recorder score was calculated for each spacer pairing as the product of four independently measured events: efficiency of pegRNA_A◊B_ targeting to the recording locus, efficiency of message+spacer B insertion by prime editing, efficiency of pegRNA_B◊A_ targeting to the recording locus, and efficiency of message+spacer A insertion by prime editing (Figure 2g). The ARCHIVE recorder (reciprocating between spacer ARCH20 and ARCH36) exhibited a ∼20-fold increase in iterative recording efficiency compared to the peCHYRON control (Figure 2h).

### Design and Quantification of the Optimized ARCHIVE System

ARCHIVE pegRNA expression architecture enabled both simplification of recorder design and increased sequential recording efficiencies (Figure 3a). PeCHYRON employs a nicking sgRNA (ngRNA), a third RNA species alongside the two pegRNAs, to suppress off-target editing of pegRNA expression vectors. The pegRNA_A◊B_ cassette may nonetheless remain susceptible to PE-complexed pegRNA_B◊A_ binding or nicking of the pegRNA-encoding strand – and vice versa – which can reduce pegRNA expression. ARCHIVE addresses this at the cassette level (below) and additionally transitions pegRNA expression from a Pol-III (hU6, H1) to a Pol-II promoter system, opening access to a wider range of constitutive (e.g., CMV) and inducible (e.g., small-molecule-, enhancer-, or response-element-driven) input modalities. Use of a Pol-II-based expression system further permitted installation of small high-splicing-efficiency introns within the RTT-encoded spacer, broadly cloaking pegRNA expression cassettes from PE recognition and consequent off-target editing, whilst reconstituting intact and functional pegRNA at the transcript level. Additionally, implementation of a Csy4-based pegRNA excision mechanism permitted use of a single promoter input to drive bicistronic expression of pegRNA_A◊B_ and pegRNA_B◊A_ simultaneously.

**Figure 3:**
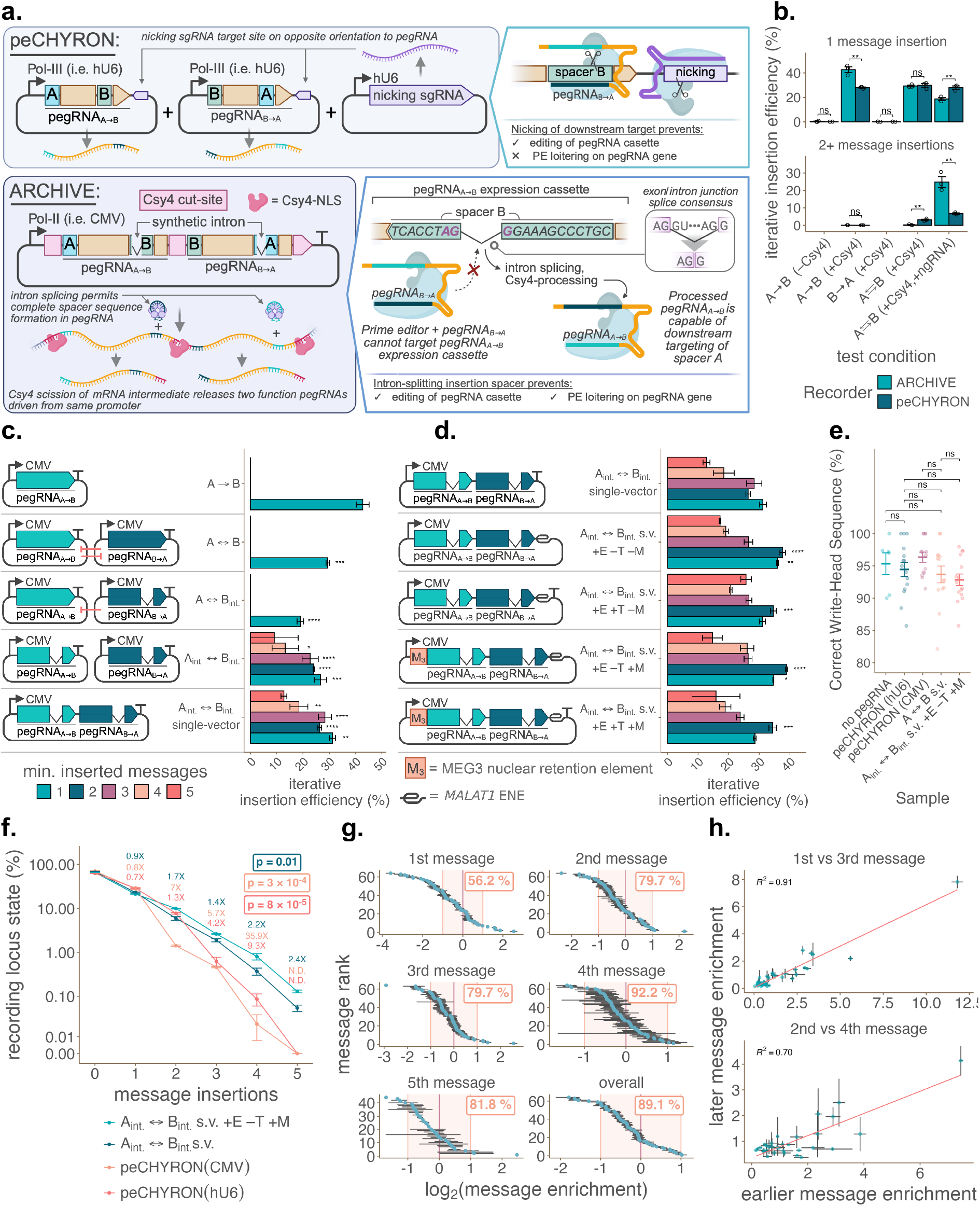
Robust Sequential Integration of Messages with ARCHIVE. **(a)** Schematic of ARCHIVE and peCHYRON systems. peCHYRON leverages use of two separate hU6-based pegRNA expression vectors complemented by a nicking sgRNA (ngRNA) to reduce unwanted pegRNA vector editing. ARCHIVE expresses both pegRNAs from a single bicistronic CMV-based expression vector, with introns inserted within RTT-encoded spacer sequences to mitigate off-target pegRNA cassette editing and PE obstruction of pegRNA expression. **(b)** Comparison of iterative insertion efficiency (number of reads with >n insertions over number of reads with ≥n insertions) for 1st and ≥2nd message recordings across test conditions for both peCHYRON and ARCHIVE. All pegRNAs were expressed from CMV-driven vectors. Status of a pEF-1α-Csy4 and ngRNA plasmid supplementation during transfection is indicated for each test condition. Test condition arrows denote pegRNA target/insertion sequence directionality, with bidirectional markers representing both A◊B and B◊A pegRNA co-transfection. Significance was calculated with two-way ANOVA with post-hoc Tukey HSD, n = 3 biological replicates. **(c)** Iterative insertion efficiency of segregated, single-vector, and intron-split pegRNA expression vectors (depicted diagrammatically) stratified by record length. Red inhibition symbols denote where a non-intron-split pegRNA are predicted to inhibit its corresponding pegRNA partner. ‘int.’ sample subscripts denote pegRNAs which contain an intron within the RTT sequence. Significance was calculated using Dunnett’s test against the pegRNAA◊B control, n = 3 biological replicates. **(d)** Iterative insertion efficiency of intron-split single-vector pegRNA expression vectors containing additional elements to enhance nuclear retention (depicted diagrammatically) stratified by record length. ‘s.v.’ denotes a single-vector pegRNA design. ‘E’ = element of nuclear expression (ENE), ‘T’ = SV40 terminator, ‘M’ = MEG3 nuclear retention element (NRE). Significance was calculated using Dunnett’s test against the pegRNAA⇄B single-vector control, n = 3 biological replicates. **(e)** Percentage of sequencing reads where the recorder write-head (the most recent spacer sequence in the record) is targetable (correct sequence identity), stratified by sample. Means are depicted as horizontal thresholds, with individual points representing fidelity readings for each record length across n = 3 biological replicates. Statistical significance was calculated using a two-way ANOVA with post-hoc Tukey HSD, with comparisons depicted for all samples against both peCHYRON (hU6-driven) and ARCHIVE pegRNAA⇄B single-vector designs. **(f)** Recording locus state frequencies for selected recorders: peCHYRON (hU6-driven), peCHYRON (CMV-driven), ARCHIVE intron-split single-vector design (Aint. ⇄ Bint. s.v.), ARCHIVE intron-split single-vector design with ENE+MEG3 elements omitting an SV40 terminator (Aint. ⇄ Bint. s.v. +E –T +M). For each record length, fold-improvement of the Aint. ⇄ Bint. s.v. +E –T +M recorder over each other sample is depicted by colour. Significance was calculated by a comparison of iterative insertion efficiencies for each recorder to the Aint. ⇄ Bint. s.v. +E –T +M sample using a one-sided t-test paired according to record length. Significance was BH-corrected and overlayed on the graph. **(g)** Log2-transformed trinucleotide message integration rates compared to the mean integration rate stratified record length for the ARCHIVE system. Shaded orange region represents a single log2-fold range from the mean integration rate, with percentage of trinucleotide messages within this range overlayed. **(h)** Comparison of message integration rate enrichment versus mean integration rates across record lengths. Correlation between ARCH-20-targeted integrations (1st versus 3rd integrated message) and ARCH-36-targeted integrations (2nd versus 4th integrated message) are depicted, in addition to R2 value and line of best fit. In all cases, error bars represent S.E.M. Statistical significance: **** p < 0.0001, *** p < 0.001, ** p < 0.01, * p < 0.05.

Initial quantification of CMV-driven recording with both peCHYRON and ARCHIVE using single genomic-copy landing-pad-harboured recording loci confirmed that (1) lack of Csy4 expression abrogated editing; (2) the ARCHIVE pegRNA_A◊B_ efficiency was modestly (∼1.5×) higher than peCHYRON_A◊B_; (3) in the absence of intron-placement within pegRNAs, the ngRNA was required for high-efficiency multi-message recording; (4) peCHYRON retained a higher percentage of reads harbouring a single message recording; (5) ARCHIVE exhibited multi-fold increases in multi-message record frequency (∼3.6×; Figure 3b). An iterative insertion efficiency metric was defined as the cumulative conditional probability of additional insertions given the presence of previous insertions, i.e. *P(>n | ≥n)* for a given number of *n* insertions.

### Optimizing Sequence-Determinants of Polycistronic pegRNA Cassettes

Initial testing of sequential ARCHIVE recording revealed that, whereas pegRNA_A◊B_ alone achieved ∼45% single-message integration by six days post-transfection, co-expression of pegRNA_A◊B_ and pegRNA_B◊A_ resulted in mutual inhibition of pegRNA expression and reduced first-message integration (∼30%), with negligible recording of additional messages (<2%; Figure 3c). Additionally, substituting pegRNA_B◊A_ with an intron-split RTT variant further reduced first-message integration to 19%, the lack of pegRNA_B◊A_ suppression precipitating increased inhibition of pegRNA_A◊B_. Efficient recovery of multi-message reads was possible only when both pegRNAs were intron-split. Placement of both pegRNAs within a single bicistronic cassette maintained recording efficiency.

To mitigate nuclear export of the bicistronic pegRNA transcript–potentially interfering with scission by nuclear-localized Csy4 and subsequent PE complexation–pegRNA expression vectors incorporating the MALAT1 expression and nuclear-retention element (ENE) and the MEG3 nuclear-retention element (NRE) were combinatorially assessed (Figure 3d) ^34,35^. Furthermore, vectors which substituted the SV40 terminator with ENE in the canonical intron-split pegRNA vector were tested, as a canonical terminator can be dispensable in this case^36^. Use of ENE without an SV40 terminator significantly increased cumulative 1^st^ and 2^nd^ message iterative integration efficiencies when coupled with the MEG3 NRE upstream of the bicistronic pegRNA cassette (1.1X and 1.48X respectively). Furthermore, use of a spliced pegRNA did not significantly reduce the sequence fidelity of the recorder’s write-head (the most recent spacer; –0.1%), thereby permitting subsequent recording on ∼93% of active loci (Figure 3e).

### ARCHIVE Demonstrates Open-Ended Recording Efficiency Improvement

Over a six-day time course, the ARCHIVE ENE+MEG3 design achieved ∼9-fold (vs hU6-driven peCHYRON) and ∼36-fold (vs CMV-driven peCHYRON) improvements in four-message record frequency (Figure 3f). CMV architecture represented a realistic ‘on’ state for Pol-II-driven inducible recording. Furthermore, ARCHIVE produced a detectable population of cells harbouring five recorded messages (0.13% of all reads, ∼15% of 4-message reads), which were absent in both peCHYRON conditions across all replicates.

### ARCHIVE Allows Substantial Variety in Message Signatures

ARCHIVE permitted efficient recording across a variety of trinucleotide message signatures. Averaged across record lengths, 89% of trinucleotide messages were integrated within one log2 unit of the mean integration rate (Figure 3g). Furthermore, message-integration efficiencies relative to the mean remained consistent across record lengths (R^2^ = 0.91 for 1^st^ vs. 3^rd^ message, inserted at ARCH-20; R^2^ = 0.70 for 2^nd^ vs. 4^th^ message, inserted at ARCH-36; Figure 3h).

## Discussion

This work builds on the peCHYRON platform, coupling ML-guided pegRNA design with experimental validation to increase iterative recording efficiency. ARCHIVE yields a ∼20-fold increase in iterative recording efficiency beyond the peCHYRON control and ∼9-fold (vs hU6-driven) to ∼36-fold (vs CMV-driven) gains in four-message record frequency. Notably, ARCHIVE produced a detectable and useable five-message population that was absent from previous design implementations, unlocking previously-inaccessible depth in recording resolution. This result represents a recorder able to capture more messages within the same timeframe, providing further insight into the dynamics of a biological input being tracked.

The core advance involves the repurposing of predictive models of prime editing efficiency for sequence generation (Figure 1). We chose a genetic algorithm to navigate the multi-objective sequence space of pegRNA design, over other approaches (e.g. gradient-based deep learning, simulated annealing, Bayesian optimization) that may offer less interpretability or fine-grained control over multi-objective constraints. Validation on generated sequences revealed that MinsePIE predicted insertion efficiency well for RTT-HA lengths ≤ 38 nt, whereas PRIDICT correlated poorly with experimental results. This likely reflects differences in the training data underlying each model (e.g. insertion-length ranges and target-site biases) and supports our validation-in-the-loop approach (Figure 2b-d, Supplementary Figure 1).

A core assumption implemented in this work throughout Figure 2 is that pegRNA spacer/PBS and insertion sequence efficiencies are independent (i.e. an optimal spacer/PBS pair is interoperable with an optimized RTT). Ever-decreasing DNA oligonucleotide library synthesis costs may enable higher-resolution mapping (i.e. smaller RTT-HA length windows, additional spacer sequences, PBS length ranges) of the reciprocating-pegRNA design space, and may test additional loci beyond *HEK3* as an optimal background for such designs. Confirming the portability of ARCHIVE by implementation in biologically and therapeutically-relevant cell types (e.g. hiPSCs, T-cells) would be a useful area of future work.

The requirement for two reciprocating pegRNA molecules to be simultaneously expressed for each unique input does limit scalability. As such, ARCHIVE’s adoption of ENGRAM’s Csy4-based approach to driving pegRNA expression from Pol-II promoters doubles as a reliable strategy for bicistronic pegRNA expression (Figure 3a). A structural feature of both peCHYRON and ARCHIVE is that pegRNA spacers used for sequential recording are also encoded within RTT sequences; this cognate sequence can act as an off-target site that may reduce recording fidelity over extended timescales – which we saw as an opportunity for further optimization. PeCHYRON presents an elegant solution in the form of an ngRNA, which we found to be indispensable for sequential integration for both systems (Figure 3b). We reasoned that use of a Pol-II based architecture could additionally permit use of synthetic short introns–strategically placed within RTT-encoded spacers–to mitigate off-target recording, simplify the design under one promoter, and further boost recording efficiency. Intron placement materially improved iterative recording efficiency (from negligible record detection of length ≥2 to 10–30% iterative recording efficiency across record lengths; Figure 3c,d).Recording efficiency was further enhanced by ∼2X (across record lengths) with nuclear-retention element placement within the pre-spliced pegRNA cassette (Figure 3f). This intron-cloaking strategy may be transferable to ENGRAM-based architectures for the same design reasons.

Beyond efficiency, ARCHIVE supports a broad set of distinguishable message signatures: 89% of trinucleotide messages integrated within one log_2_ unit of the mean rate, and relative integration efficiencies remained stable across record positions (R2 = 0.91 for 1st vs 3rd, 0.70 for 2nd vs 4th). This position-independence is what renders ARCHIVE usable as a programmable, multiplexed input-encoding substrate, since a fixed message-to-input mapping can be decoded reliably regardless of where in the record it appears. Future advances of DNA typewriter/ENGRAM (such as additional sgRNA scaffolds and recorder-associated parts) would benefit and complement future ARCHIVE implementations^37^.

Critically, because ARCHIVE drives pegRNA expression from a Pol-II promoter, it is in principle compatible with the inducible, enhancer-, and response-element-driven inputs that govern the majority of endogenous gene-regulatory programs–a regime structurally inaccessible to Pol-III-locked designs. We emphasize, however, that the present work establishes this capability at the level of architecture and constitutive (CMV) on-state performance; direct recording from a live inducible biological trigger was not tested here. Demonstrating faithful recording of a small-molecule-gated input, endogenous CRE activity (akin to ENGRAM implementations), or a synthetic cell-state-responsive promoter is therefore the most immediate priority for future work. Such an implementation may involve ENGRAM’s pre-validated inputs (i.e. Tet-On, NF-κB, Wnt), but could scale to recording synthetic TFBS reporter activity across experimental trajectories (e.g. stem cell differentiation).

Longevity and durability of ARCHIVE recorders stably integrated into human cells over months-long experimental trajectories remains a key area for future investigation–while initial results regarding write-head fidelity are indeed promising (Figure 3e). Finally, the relationship between ARCHIVE-recorded event ordinality and event magnitude remains to be determined–the implementation of a kinetic model to this end may enable deconvolution of analog signal intensity from DNA records. Overall, ARCHIVE fundamentally increases the efficiency of open-ended recording systems. We anticipate that it will enable deeper, non-destructive insight into cell behavior across extended timescales.

## Methods

### Machine-Guided *in Silico* Evolution of Spacers to Permit Simultaneous On-Target Prime Editor Targeting and Edit Integration

To design the first spacer library (Library 1), comprising 20-nt spacers that could be efficiently integrated into the HEK3 locus while maintaining high predicted Cas9 on-target activity, we performed the following steps: (1) generated a diverse set of candidate spacers as seeds for a genetic algorithm (GA); (2) ran the GA using MinsePIE as a surrogate model of prime editing-mediated insertion efficiency at HEK3, with MinsePIE predictions serving as the fitness function and DeepSpCas9 and RS3 scores used to threshold Cas9 on-target activity; (3) scored the off-target potential of evolved spacers using CRISPRitz-based prediction; (4) further evaluated spacer-mediated insertion efficiency using PRIDICT; and (5) selected top-scoring spacers for inclusion in Library 1 for experimental validation. Modeling was conducted in R.

### Generation of Spacers for Seeding an Island-Based SAGA

Candidate spacers for the first library were generated by random sampling of sequences with the consensus *GNNNNNNNNNNNNNNNNTGA*, with a 5′ G to ensure hU6-compatible transcription and a TGA at positions 18–20 corresponding to the 5′ end of the HEK3 RTT. Spacers were required to maintain a GC content between 40% and 60% and to contain the AGG exonic splice-junction consensus, encoded in the spacer as the reverse-complement CTT motif. For each sampled spacer, DeepSpCas9 ^38^ scores were computed locally using the published implementation (https://github.com/MyungjaeSong/Paired-Library); only spacers in the top 5% were retained. To maximize diversity, redundant spacers (pairwise Hamming distance ≤ 5) were removed, and sampling was iterated, comparing each new candidate against the growing pool, until 50,000 unique spacers were obtained.

### MinsePIE Model Tuning for High-Throughput Prime Editing Efficiency Prediction

Prime-editing insertion efficiencies were predicted using MinsePIE ^32^, which was installed in a Miniforge3 virtual environment (conda version 25.7.0, python version 3.12.8, macOS 15.2, osx-arm64 platform) on Apple M1 hardware. To enable high-throughput scoring of pegRNA designs, we modified the MinsePIE predictor to accept a CSV of pegRNA features, pass the full table to *`minsepie.predict`* in batch mode, and return insertion scores and features as a CSV. The underlying MinsePIE model (version 3.0) and feature calculations were not modified. The high-throughput-optimized fork of MinsePIE, termed MinsePIE-HT (commit #8f4624c), is available at https://github.com/aarhar12/MinsePIE-HT.

### MinsePIE-Based Surrogate-Assisted Genetic Algorithm for Iterative Evolution of Spacer Sequences

In the first round of evolution, generated spacers were filtered to remove sequences containing homopolymer runs (T_4_, A_4_, G_4_, or C_4_), after which all spacers meeting the specified sequence-composition constraints and retaining a DeepSpCas9 score of ≥60 and an RS3 ^39^ (https://github.com/gpp-rnd/rs3) score of ≥1 were scored using MinsePIE. DeepSpCas9 and RS3 on-target efficiency predictions were made on spacers of the form TTGG-spacer-TGGCAG to reflect local *HEK3* locus genomic sequence context. Spacers were then scored using MinsePIE-HT, with the canonical HEK3 targeting spacer used to insert each candidate spacer alongside a 13-nt PBS and 34-nt RTT-HA–matching the parameters used for the *HEK3*+6xHis integration. The top 1500 candidate spacers were selected to seed an identical number of isolated islands individually for 12 subsequent rounds of evolution. At the conclusion of each generation, all single-substitution variants were generated from the spacer which maximized MinsePIE z-score, and were used at equal frequency to seed the island for subsequent rounds of evolution.

In subsequent generations of evolution, spacers within each island were re-evaluated using the same DeepSpCas9, RS3, and MinsePIE-HT scoring framework described above. From generation 4 onward, spacers were required to encode a CCT-containing PAM-proximal motif, corresponding to the reverse complement of the AGG exonic splice-site consensus. This strategy permitted broader exploration of sequence space during the early generations of evolution while filtering for spacers containing the exonic splice-site consensus during later generations. Placement of the CCT motif proximal to the PAM ensured that, during downstream ARCHIVE recorder development, intron placement within the pegRNA cassette would obfuscate off-target prime editor targeting.

### Scoring Spacer Off-Target Potential Within the Human Genome

Off-target liability for candidate spacers was estimated using CRISPRitz^40^ (https://github.com/pinellolab/CRISPRitz; v2.6.6), installed using Docker. Candidate spacers were searched against the human hg38 genome, with 3 total mismatches permitted, and off-target activity was quantified using cutting-frequency-determination (CFD) scoring via *`crisprScore::getCFDScores()`* in R. For mismatch-only off-target prediction, each spacer was assigned an off-target score (OTS), defined as 1 − max(CFD) across all detected genomic off-target sites, such that higher values indicate greater predicted specificity.

To account for bulge-containing off-targets, an additional CRISPRitz search was performed allowing 4 mismatches, 1 DNA bulge, and 1 RNA bulge. Bulged alignments were processed to restore the aligned spacer/DNA representation before CFD scoring, and separate specificity metrics were calculated for bulge-free and bulge-containing off-target classes. In this implementation, OTS denotes the mismatch-only specificity score, and OTS_bulge denotes the specificity score for single-bulge off-targets. Final candidate spacers were required to exceed an OTS threshold of 0.8 and an OTS_bulge threshold of 0.4 before inclusion in the optimized spacer set.

### Selection of Top-Performing Spacer Sequences for Experimental Validation

While the evolution scheme fixed the RTT-HA length at 34 nt, spacers which passed off-target risk assessment were rescored with MinsePIE-HT across a variety of RTT-HA lengths (26–50 nt in 4-nt increments) to explore if modification of this parameter improved predicted z-score. Spacer-HA pairs that exceeded a MinsePIE z-score threshold of 0.75 (the model’s native standardized output) were additionally scored using a local command-line implementation of PRIDICT ^33^, minimally modified from the original GitHub repository (https://github.com/uzh-dqbm-cmi/PRIDICT; v1.0) by restricting the default PBS and RT search ranges in *`pridict_pegRNA_design.py`* to a *`PBSlengthrange`* of 13 nt and an *`RToverhanglengthrange`* of 26–50 nt in 4-nt increments. Top spacer sequences were selected by maximizing the MinsePIE z-score within each of three cohorts: (1) high off-target specificity, defined as the 90th percentile of PRIDICT scores with OTS > 0.9 and OTS_bulge > 0.5; (2) relaxed off-target specificity, defined as the 90th percentile of PRIDICT scores with OTS > 0.8 and OTS_bulge > 0.4; and (3) relaxed off-target specificity without PRIDICT-based resolution, defined by OTS > 0.8 and OTS_bulge > 0.4. These thresholds were chosen heuristically to sample a range of performance– specificity trade-offs.

### Cloning of pegRNA Libraries for Spacer Integration Efficiency Testing

Spacers were ordered as an oPool oligo library (Integrated DNA Technologies, Coralville, IA, USA), with each element conforming to the consensus sequence *ACAGGTCTCCA– (spacer reverse complement)–NNN–CGTGTGAGACCTATTCGCC*, where the degenerate trinucleotide represents all possible messages to be recorded. Library elements were amplified by PCR with Q5 Hot Start High-Fidelity 2X Master Mix (New England Biolabs / NEB, Ipswich, MA, USA; M0494). In addition, both peCHYRON–A/B spacers were included as controls. A *HEK3-*targeting U6-driven pegRNA expression plasmid derived from pU6-tevopreq1-GG-acceptor (Addgene plasmid #174038) was used as a backbone template. The oligo library was included as a forward primer binding the PBS, while the *pR4Dest_BB_R* reverse primer targeted the end of the pegRNA scaffold. RTT-HA elements of varying length (26–50 nt, in 4-nt increments) were synthesized by Primerize-based PCR assembly using primers *RT_xx_F/R*, where xx denotes the RTT-HA length^41^. For each reaction, 1 μL of each primer (100 μM) was combined with 12.5 μL Q5 Hot Start High-Fidelity 2X Master Mix and brought to 25 μL with nuclease-free water, then cycled 23 times under the manufacturer’s standard conditions. PCR samples were purified using the Monarch® Spin PCR and DNA Cleanup Kit (NEB; T1130). Plasmid libraries (corresponding to each RTT-HA length) were constructed by Golden Gate assembly using library-tagged vector backbone and each RTT element as an insert using NEBridge Golden Gate Assembly Kit (BsaI-HF v2; NEB; E1601) according to the manufacturer’s protocol. Assemblies were purified with PCR cleanup columns, followed by transformation using NEB Stable chemically-competent *E. coli* (NEB; C3040) according to standard protocols. Samples were plated on LB-agar (BioShop, Burlington, ON, Canada; LBA408) with 100 μg/mL carbenicillin (BioShop; CAR666) and incubated overnight at 30 °C before plasmid isolation. For each plate, colonies were homogenized by scraping with a plate-spreader submerged in 1 mL of ice-cold LB-broth, prior to plasmid preparation using the PureLink Quick Plasmid Miniprep Kit (Invitrogen, Waltham, MA, USA; K210010). Library coverage for each RTT-HA length exceeded nominal depth (≥100× coverage; 121 elements), assessed by colony-counting on serial dilutions of a pooled transformation mixture (across RTT length-specific libraries). Plasmid libraries were sent for Sanger sequencing at Eurofins Genomics to confirm pegRNA sequence fidelity and library sequence degeneracy using the hU6F sequencing primer. Finally, a pegRNA vector encoding a HEK3 6×His-tag insertion was spiked in at ∼1:100 (mass) as an internal control, representing a pre-validated high-efficiency large insertion at HEK3. Vectors were labelled as pU6-R4-pegRNALib_RT*xx*, where *xx* denotes the RTT-HA length. All plasmids used in this study (excluding pegRNA constructs used in Figure 3) are catalogued in Supplementary Table 2.

### General Human Cell Culture Methods

HEK293T cells (thawed at passage 7) were routinely cultured in DMEM (Gibco, Waltham, MA, USA; 11965092) supplemented with 10% fetal bovine serum (FBS, Value-grade; Gibco; A5256701), and subcultured every 3-4 days by dissociation with TrypLE (Gibco, 12605028) followed by standard cell culture methodologies: Centrifugation at 500 rcf for 5 mins, supernatant aspiration, resuspension in fresh medium, quantification using a Countess 3 automated cell counter (Invitrogen; AMQAX2000), and plating at a desired density. Cells were maintained at 37 °C and 5% CO2 under ambient O_2_ in a humidified incubator. Cells were routinely transfected at 60%+ confluency using Lipofectamine 3000 Reagent (Invitrogen; L3000015) according to the manufacturer’s protocol using Opti-MEM (Gibco; 31985062) medium. When necessary, cells were cryopreserved in complete medium with 10% DMSO (Thermo Scientific Chemicals, Waltham, MA, USA; J666650.AP) and frozen at a controlled rate in a Mr. Frosty container (Thermo Scientific, Waltham, MA, USA; 5100-0001) in a −80 °C freezer, followed by long-term banking under liquid nitrogen. Cells were routinely assessed for morphology and density using an ECHO Revolve Microscope (ECHO, San Diego, CA, USA). Cells were routinely tested to be negative for mycoplasma contamination^42^.

### Experimental Validation of Spacer Integration in the *HEK3* locus

#### Laboratory Methods

HEK293T cells were seeded at 60% confluency in seven T-25 flasks. The next day, each flask was co-transfected with one of the seven RTT-length-matched pegRNA libraries and the pCMV-PE2-P2A-EGFP prime editor–EGFP vector (library:editor 1:3, mass). Cells were incubated for 3 days before fluorescence-activated cell sorting (FACS). In brief, cells were dissociated, washed with phosphate buffered saline (PBS; Wisent Bioproducts, Saint-Jean-Baptiste, QC, Canada; 311-010), and each sample was stained with LIVE/DEAD Fixable Near-IR viability dye (Invitrogen; L10119; 1:500 in 1 mL PBS) for 30 min in the dark. Cells were then washed with PBS and resuspended in FACS Buffer consisting of PBS supplemented with 1% bovine serum albumin (BSA; Wisent Bioproducts), before sorting on a BD FACSAria IIIu (BD Biosciences; 809-098). A representative gating strategy, outlining cell gating, singlet isolation, viability gating, and GFP gating is depicted in Supplementary Figure 1f.

Sorted cells were centrifuged at 800 rcf for 5 min followed by aspiration of the supernatant. Cell pellets were resuspended in 200 μL of PBS followed by gDNA isolation using the PureLink Genomic DNA Mini Kit (Invitrogen; K182001). For each RTT-HA length-specific library, PCR of the *HEK3* genomic locus was conducted using the entire gDNA sample using Q5 Hot Start High Fidelity 2× Master Mix for 30 cycles (parameters optimized to minimize overamplification) using *HEK3_NGS_F/R* primers containing Amplicon-EZ-compatible (GENEWIZ/Azenta Life Sciences) adapter sequences, where the forward primer contained an additional 7-nt index sequence to denote RTT-HA length-specific library identity. Similarly, the RTT-region of each pegRNA library was amplified with *R4_PL_NGS_F/R* primers. PCR products were purified prior to pooling and submission to GENEWIZ (Amplicon-EZ; Azenta Life Sciences, South Plainfield, NJ, USA) for Illumina sequencing.

#### Computational Analysis Pipeline

Sequence data was read into R using the *ShortRead* package, and plasmid library reads were separated from those corresponding *HEK3* loci. RTT-HA lengths were determined for plasmid library-specific reads, and relative abundance of each spacer, and its encoded trinucleotide message was calculated. Similarly, RTT-HA lengths were associated with *HEK3*-locus-specific reads based on index barcode identity, and relative abundance of each spacer, its encoded trinucleotide message, and RTT-HA length was calculated. Enrichment scores for each spacer were determined based on the ratio of *HEK3* locus edit abundance versus plasmid library abundance for the same RTT-HA length and spacer, with further stratification based on trinucleotide message identity. A reconstructed editing efficiency score was determined by scaling the enrichment score by overall *HEK3* locus editing efficiency, weighted by plasmid library spacer abundance for each RTT-HA length as follows:

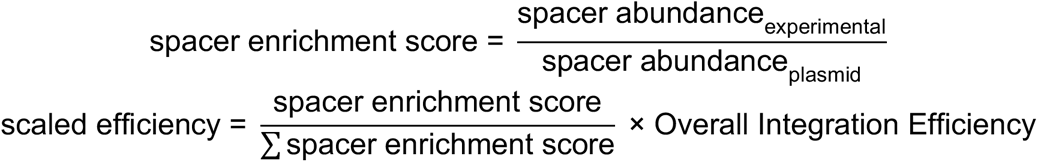

Per-spacer standard error, weighted by trinucleotide message abundance, was computed with *diagis::weighted_se()* in R.

### Repetition of Spacer Evolution and Validation Using a Random Forest Model Trained on Initial Experimental Data

To improve the insertion-efficiency fitness model beyond MinsePIE z-score alone, a random forest was trained on the initial library dataset. Only insertion data from RTT-HA lengths ≤ 38 nt were used, owing to MinsePIE’s poor predictive power at RTT-HA lengths ≥ 42 nt. RTT MFE, MinsePIE z-score, spacer position-15 nucleotide, RTT-HA length, and PRIDICT on-target score were used as predictive features. The model used 350 trees and mtry = 2, evaluated by 5-fold cross-validation repeated 10× with folds stratified by spacer identity to prevent leakage across RTT-HA length replicates of the same spacer. The final model achieved a mean cross-validated R² of 0.51 ± 0.06 across held-out spacers, compared to a training R² of 0.70, indicating moderate overfitting consistent with the limited training set size (n = 116 spacers). Model feature importance was assessed using Shapley additive explanations (SHAP) values computed in R with *`fastshap::explain()`* using a 100-simulation Monte Carlo approximation, confirming RTT MFE and MinsePIE z-score as the most influential predictors of insertion efficiency. Spacer evolution was repeated as previously described, with differential selection criteria following feature prediction steps. Firstly, an additional CFD_intron_ score was calculated based on the permissiveness of a given spacer for an intron-split target site. CFDintron was defined as the maximum CFD score across the 20 PAM-proximal nucleotides generated by inserting each of five synthetic intron splice-acceptor sites (Supplementary Table 4) into the exonic splice consensus present on each spacer. As such, this score was a rough estimate of intron-based cloaking of pegRNA expression cassettes from off-target pegRNA targeting. All spacers exhibited a CFD_intron_ score of ≤0.05, and an OTS_bulge score of ≥0.6. To reduce sequence redundancy and broaden the search, spacers were hierarchically clustered on global pairwise-alignment scores (*`pwalign::pairwiseAlignment`*). The 116 spacers maximizing predicted efficiency or MinsePIE z-score were selected for testing as for the initial library. Library 2 vectors were labelled as pU6-R5-pegRNALib_RT*xx*, where *xx* denotes the RTT-HA length.

### Library Design for Spacer Targeting Across a Fixed set of Defined Spacer-length Edits

Top-performing spacers for HEK3 integration (16 from library 1, 25 from library 2, plus peCHYRON-A/B controls) were selected as prime-editing targets, using a fixed set of six spacer-length insertion edits (also drawn from library 2). Each spacer for testing was ordered as an oligonucleotide library, whereby each element consisted of the spacer, inverted BsmBI cut sites (for downstream Golden Gate cloning), a PBS (of length 11, 13, or 15) in addition to a target site flanked by BbsI cut sites and PCR handles. As such, each library element encoded sequence information for both a pegRNA expression cassette and its corresponding target site on a single topology. Unless otherwise stated, all PCRs in this section used Platinum SuperFi II Green PCR Master Mix (Invitrogen; 12369010) in 25 μL reactions per the manufacturer’s protocol. The library was amplified for 12 cycles using *R5L2_S1_F/R* primers. Additionally, a FAST-STEM pGLP2_PuroR backbone (Addgene #259078) carrying the hU6 promoter and HEK3 RT homology arm was linearized by PCR with *R5L2_BB_F/R* (encoding BbsI sites complementary to the amplified library), then treated with 10 U DpnI (NEB; R1076) at 37 °C for 1 h. Following PCR-purification of both insert and backbone PCRs, assembly was conducted by Golden Gate using BbsI-HF (NEB; R3539M) and NEBridge Ligase Master Mix (NEB; M1100) according to the manufacturer’s protocol in a 40 μL reaction, which was subsequently PCR-purified. The purified assembly was electroporated into in-house electrocompetent NEB Stable *E. coli* in four reactions (25 μL cells per reaction; 1-mm cuvette) using a BTX Gemini X2 (Harvard Apparatus, Holliston, MA, USA; 45-2040) using the following settings: 2.0 kV, 200 Ω, and 25 μF. Outgrowth was conducted for 90 min at 30 °C in NEB 10-beta/Stable Outgrowth Medium (NEB; B9035S) followed by plating on LB+Agar supplemented with 100 μg/mL carbenicillin overnight at 30 °C. Plasmid preparation from solid medium was conducted the following day as previously described. This vector library is hereafter referred to as ‘level-1’ in subsequent uses.

The level-2 vector library was constructed by integration of a pegRNA-scaffold/RTT/spacer oligo pool into the level-1 backbone library. The oligo pool was constructed by *Primerize*-based assembly using *R7_scaffold_B1/B2/B3/B4* primers, where the B4 primer was a small oligo pool encoding all six test spacers for insertion (R7_scaffold_B4A–F). Construction of the level-2 vector was conducted by Golden Gate assembly using the NEBridge Golden Gate Assembly Kit (BsmBI; NEB; E1602) according to the manufacturer’s protocol in a 40 μL reaction. Assembly reaction purification, electroporation, outgrowth, expansion, and preparation were conducted as described for the level-1 vector library.

Finally, the level-3 vector library was constructed by PCR amplification of the level-2 library backbone using *R7_BB_F/R* primers which additionally encoded PaqCI cut-sites. This PCR linearized the level-2 vector library between the pegRNA and target-site segments. A UMI sequence insert was assembled by Primerize (as previously described) using *R7_UMI_F/R* primers. Finally, Golden Gate assembly was used to integrate UMI elements into the linearized level-2 vector backbone using PaqCI (NEB; R0745) and T4 DNA Ligase (NEB; M0202) according to their supplied protocol. Assembly reaction purification, electroporation, outgrowth, expansion, and preparation were conducted as described for the level-1 vector library.

### Derivation of a HEK293T Landing Pad Cell Line

HEK293T cells were thawed, and a landing pad encoding the EF-1α promoter driving Neomycin resistance marker (NeoR) and EGFP transgenes (NR landing pad) was engineered into the *AAVS1* locus at single-copy in an identical manner to Rosenstein *et al.*^43^ with the following modifications: (1) Lipofectamine 3000 was used instead of Lipofectamine Stem; (2) A plasmid donor was used instead of lssDNA; (3) Monoclonal expansion used limiting dilution at 0.5 cells/well in 96-well plates; (4) the ddPCR copy-number ratio for single-copy insertion was ∼0.66, consistent with the triploid RNaseP reference in HEK293T (1 copy NeoR / 3 copies RNaseP = 0.66). The selected clone is hereafter referred to as ‘HEK293T-LP’.

### Experimental Validation of Spacer Targeting Across a Fixed set of Defined Spacer-length Edits

#### Delivery of the Level-3 pegRNA-Target Site Library to 293T Landing Pad Cells and Subsequent Prime Editing Induction

HEK293T-LP cells were seeded at 70% confluency in a T-75 flask. The next day, the level-3 vector library and a Bxb1 expression vector (pBxb1; Addgene #259095) were co-transfected at 4:1 (mass) with Lipofectamine 3000 (Invitrogen), scaled per the manufacturer’s protocol. Three days later, medium was supplemented with 2 μg/mL puromycin (Millipore Sigma, Burlington, MA, USA; P7255); cells were passaged for 7 days in a T-75 flask, after which selection was stopped. Cells were then seeded at 70% confluency for transfection in a T75 flask and transfected with pCMV-PE2-P2A-BleoR plasmid with Lipofectamine 3000. The next day, 400 μg/mL Zeocin (Gibco; R25001) was added, and cells were passaged two days later. Six days post-transfection, cells were dissociated and PBS-washed, following gDNA extraction with the PureLink gDNA mini kit (one spin-column per 5 million cells).

#### Next Generation Sequencing Library Preparation

Extracted gDNA was subjected to two parallel PCRs amplifying the target-locus and pegRNA-cassette regions with *R7_RL_NGS_F/R* and *R7_pegRNA_NGS_F/R*, respectively, appending Illumina Nextera i5/i7 overhang adapters. Both amplicons were inclusive of the genomically-integrated UMI region for linking pegRNA/target locus identity. As such, PCR amplification steps did not include primers encoding UMIs. PCRs were conducted for 30 cycles with Platinum SuperFi II Green PCR Master Mix, followed by purification. A secondary 5-cycle PCR added standard Nextera 10-nt dual-index adapters; products were purified and submitted to the University of Wisconsin–Madison Biotechnology Center (UWBC) for sequencing on an Illumina NovaSeq X (2×150 PE; shared flow cell; ∼20M reads ordered).

### Computational Analysis of Spacer Targeting Across a Fixed set of Defined Spacer-length Edits

pegRNA features were extracted from raw Illumina reads and matched to target-locus reads by UMI identity in R. Only pegRNA/target pairs in which the spacer sequence, PBS, and target site were cognate were retained for analysis. For each spacer/PBS-length combination, mean editing efficiency was calculated across all six test insertion sequences. Only spacer/PBS-length combinations with a minimum of 15 readings (unique UMIs) were retained for analysis to maintain nominal precision of efficiency estimates.

To identify top recorders (two linked spacers with PBSs enabling reciprocal targeting), a recorder score was computed as the product of four efficiencies: Cas9 targeting of the first spacer (targeting_A_), insertion of the second spacer (insertion_B_), Cas9 targeting of the second spacer (targeting_B_), and insertion of the first spacer (insertion_A_). As such, recording score indicated the cumulative frequency of completing all four discrete actions in the recording loop. pegRNAs predicted to form PBS–RTT secondary structures were excluded to avoid interference with the spacer/flap region.

### Design of the Optimized ARCHIVE System

#### Cloning of pegRNA Constructs

Individual pegRNAs were cloned into either CMV or U6 promoter-encoding plasmid backbones using NEBridge Golden Gate Assembly kit (BsaI-HFv2). The nicking sgRNA spacer was identical to that used by peCHYRON, with a target site placed downstream of the pegRNA locus in both promoter classes of vector. For the CMV-based backbone, Csy4 hairpins flanked the pegRNA insertion site. For each CMV-expressed pegRNA, an 8-nt linker that minimized secondary structure with the PBS+RTT (predicted by *`ViennaRNA RNAfold`* v2.5.1) was placed between the PBS and the 3′ Csy4 site^44^. The nicking sgRNA was cloned into the pU6-pegRNA-GG-acceptor backbone (Addgene #132777) according to the protocol employed by Anzalone *et al*^45^. pegRNA fragments were generated using Primerize with primer-encoded degenerate trinucleotide message sequences. For dual pegRNA expression vectors (pegRNA_A◊B_ & pegRNA_B◊A_), BbsI was used to generate cohesive DNA ends downstream of the 3′ Csy4 site, enabling subsequent integration of an additional pegRNA with BsaI-based Golden Gate assembly. Finally, for pegRNA expression constructs containing MEG3 NREs or MALAT1 ENEs, such elements were pre-cloned into custom CMV-based pegRNA expression vectors. All constructs (U6 and CMV, peCHYRON and ARCHIVE) were assembled according to an extended Golden Gate assembly regime ((5 min at 37 °C, 5 min at 4 °C) × 60 cycles, 5 min at 60 °C). Assembly reactions were purified with solid-phase reversible immobilization (SPRI) beads (1× v/v beads:reaction)^46^ and transformed into NEB Stable *E. coli* made competent with the Mix & Go! kit (Zymo Research, Irvine, CA, USA; T3001). Cells and DNA were mixed, incubated on ice for 15 min, heat-shocked at 42 °C for 45 s, returned to ice for 5 min, and outgrown in NEB 10-beta Stable Outgrowth Medium for 30 min at 30°C. Cells were spread on LB+Carbenicillin plates and incubated overnight at 30 °C, followed by pooled colony harvesting and plasmid preparation the following day. All pegRNA constructs used in this study are listed in Supplementary Table 3.

#### Cloning and Delivery of Recording Loci to HEK293T Landing Pad Cells

Primerize PCR was employed to generate both the ARCH-20 and peCHYRON recording loci, with flanking *HEK3* locus sequences, followed by Golden Gate assembly into the FAST-STEM pGLP1_PuroR vector (Addgene #259077) using the NEBridge Golden Gate Assembly kit (BsmBI) according to the manufacturer’s protocols. Each recording-locus plasmid was delivered to HEK293T-LP cells as described above.

#### Quantification of Iterative Recording Efficiency

HEK293T-LP populations pre-engineered with recording loci were seeded at 75% confluency in 96-well plates. Transfection mixtures were performed as follows using a high-efficiency prime editor, with Csy4 and ngRNA added where indicated: (i) 250 ng pCAG-PEmax; (ii) 125 ng pEF-1α-Csy4-NLS; (iii) 125 ng of each pegRNA vector (or 250 ng for single-vector designs); (iv) 125 ng pU6-ngRNA. Reactions were prepared according to the standard 24-well Lipofectamine protocol (1.5 μL Lipofectamine 3000, 1 μL P3000 reagent, 25 μL Opti-MEM in each tube) according to the manufacturer’s protocols, with 10 μL of transfection mixture dispensed to respective wells in triplicate. Medium was refreshed the following day, and cells were expanded for 3 days prior to passaging at a 1:5 split ratio. Six days post-transfection, medium was aspirated and cells were lysed by incubation at 37 °C *in situ* with 10 mM Tris-HCl (pH 7.5; Molecular Biology Grade; Promega, Madison, WI, USA; H5123), 0.05% SDS (BioShop; SDS001.500), 25 μg/mL Proteinase K (Invitrogen, 25530049) for 2 hours followed by 80 °C for 30 min. gDNA was purified from lysates by SPRI-based cleanup at a 2× bead:lysate ratio (v/v).

Purified gDNA was PCR-amplified for five cycles with Platinum SuperFi II DNA polymerase using an N7 UMI-tagged forward primer (*R8_NGS_PCR1_UMI_F*) and one of three replicate-indexing reverse primers (*R8_NGS_PCR1_R_rep1–3*). This PCR additionally appended Illumina Nextera i5/i7 overhang adapter sequences. First-round PCRs were SPRI-purified, then indexed by a secondary PCR with standard Nextera 10-nt dual-index primers to delineate sample identity for 30 cycles. Second-round PCRs were purified, normalized, and pooled prior to submitting for Illumina sequencing at UWBC (20M nominal reads requested).

#### Analysis of ARCHIVE and peCHYRON Recorder Data

NGS reads were demultiplexed and further stratified by replicate identity, and reads containing identical UMIs (or with a UMI within a Levenshtein distance of 2 of a single preexisting UMI) were collapsed. Recorded loci were tagged with an 8-nt UMI (4⁸ = 65,536 possible sequences). Approximately half of each sample’s gDNA entered the first-round PCR, and ∼15K unique UMIs were recovered per sample replicate – roughly 20–25% of the UMI space – indicating that the UMI diversity was not saturated and that collision-driven undercounting was minimal (estimated ≲10–15%) and approximately uniform across record classes. For both peCHYRON and ARCHIVE, the following per-sample metrics were determined: (i) the frequency of each record length among deduplicated reads; (ii) the identity of each trinucleotide message within records; and (iii) the sequence fidelity of the write-head (the last spacer in the record). Record frequencies were scaled for the ∼60% transient transfection efficiency observed. Iterative recording efficiency was calculated as the frequency of a subsequent record addition given the prior pool of reads of a defined length or greater.

### Data Analysis Methods

R (v4.5.2) was used for all modelling, evolution, and analysis workflows using Rstudio (latest used version was 2025.09.2+418). R packages used include data.table (1.18.2.1), dplyr (1.2.0), ggplot2 (4.0.2), Biostrings (2.78.0), e1071 (1.7.17), pbapply (1.7.4), stringr (1.6.0), stringi^47^ (1.8.7), compiler (4.5.2), parallel (4.5.2), reticulate (1.45.0), crisprScore (1.14.0), dqrng (0.4.1), randomForest (4.7.1.2), ggthemr (1.1.0), tidyr (1.3.2), ungeviz (0.1.0), ggpubr (0.6.3), ggrepel (0.9.7), ggtext (0.1.2), pwalign (1.6.0), rstatix (0.7.3), ggpmisc (0.6.3), facefuns (0.0.0.9000), mgsub (1.7.3), broom (1.0.12), ShortRead (1.68.0)^48^, readxl (1.4.5), purrr (1.2.1)

### Inclusion and Ethics Statement

The research project described herein included local researchers in design, execution, analysis and publication authorship. The authors believe this research to be locally relevant, and has attracted interest from local partners. Authors agreed on a research plan, including involvement of local trainees prior to conducting experimentation. This research was not restricted in the setting of the researchers. No animal models were used in this work. No stigmatization or risk to researcher health, safety, or security was encountered. Citations were chosen based on relevance and appropriateness for substantiating claims made, and local and regional research was taken into account when possible.

## Acknowledgements

This work was supported by Canadian Institute for Health Research and CFREF Medicine By Design funding.

## Data and Materials Availability Statement

Data and analysis conducted in this study are present in both main-text and supplementary information and data. Next generation sequencing data generated in this study will be made available through the Sequence Read Archive (SRA). Plasmids pertaining to this work will be made available on Addgene. No plasmid constructs described in this work are proprietary or patent-pending, and all are readily available for use by the community.

## Supplementary Data

**Supplementary Figure 1:**
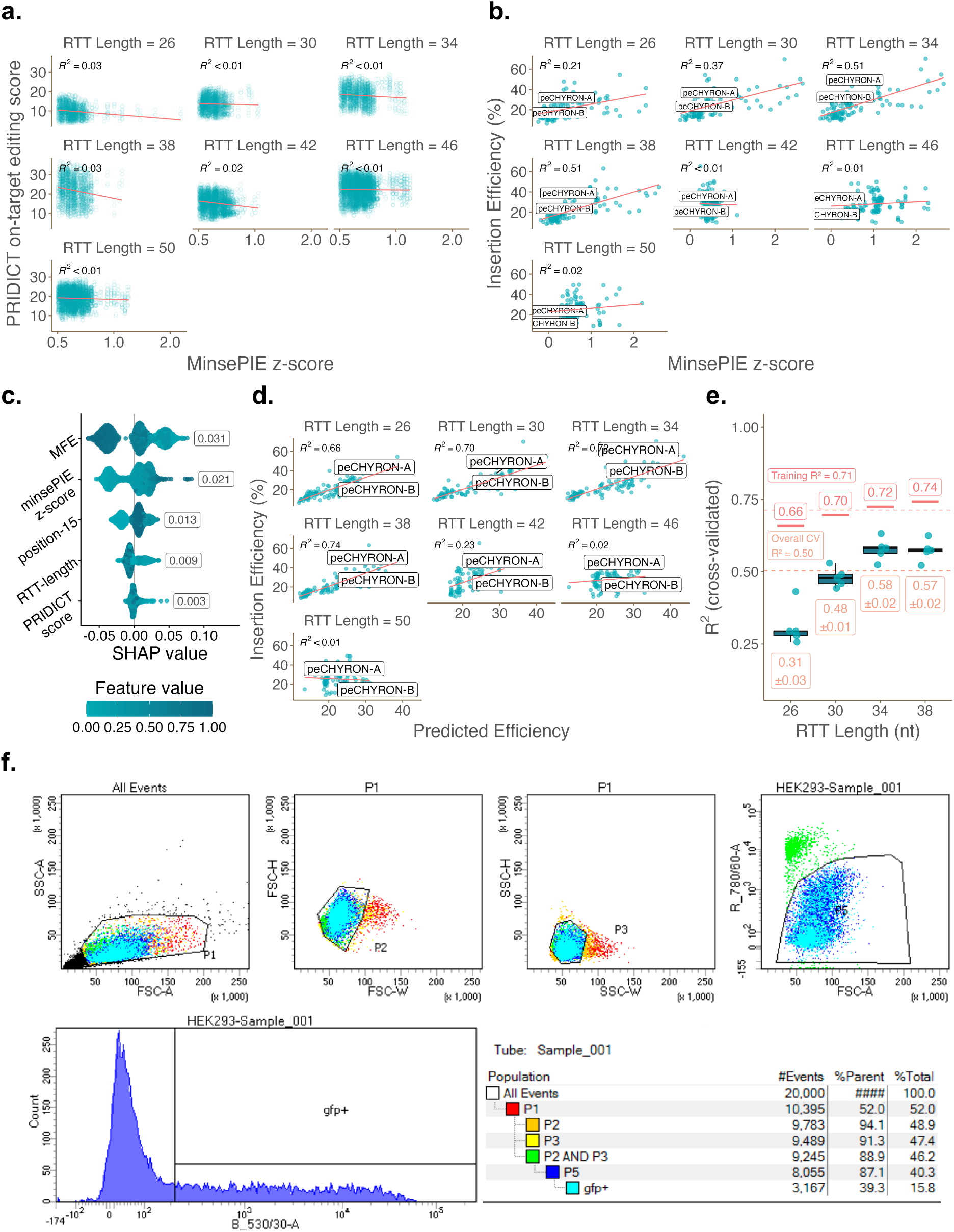
Machine-Guided *in Silico* Evolution of Spacers to Permit High-Efficiency Edit Integration, and Validation Thereof. **(a)** Correlation between MinsePIE z-score and PRIDICT on-target score across all evolved spacer sequences, stratified by RTT homology arm length. Line of best fit and R^2^ are overlayed on each plot facet. **(b)** Correlation between MinsePIE z-score and scaled insertion efficiency across all experimentally validated spacer sequences, stratified by RTT homology arm length. Line of best fit and R^2^ are overlayed on each plot facet. **(c)** SHAP beeswarm plot of random forest (RF) features (RTT MFE, MinsePIE z-score, spacer position 15 nucleotide, RTT-HA length, and PRIDICT on-target score) used to predict insertion efficiency. Mean absolute SHAP values for each feature are displayed beside each beeswarm respectively. **(d)** Correlation between RF-predicted and experimentally determined insertion efficiency across all tested spacer sequences, stratified by RTT homology arm length. Line of best fit and R^2^ are overlayed on each plot facet. **(e)** Cross-validated random forest model performance across RTT-HA lengths used for training. Boxplots display the distribution of R² values across 50 cross-validation splits (5-fold × 10 repeats), stratified by spacer identity to prevent data leakage between RT-length replicates of the same spacer. Red horizontal bars and labels indicate in-sample training R² per RTT-HA length from the full model; Orange dashed line indicates overall training and cross-validated R², respectively. Mean cross-validated R² ± S.E.M. across splits is shown below each group. **(f)** Flow cytometric gating strategy for FACS-sorting HEK293T cells transfected with pegRNA libraries and Subsequent NGS Analysis Workflow. Cells were identified by FSC-A/SSC-A gating, followed by singlet isolation (FSC-W/FSC-H; P2) → (SSC-W/SSC-H; P3) in addition to viability gating (LIVE/DEAD-Near-IR; P5), prior to gating for EGFP+ cells (gfp+).

**Supplementary Table 1:**
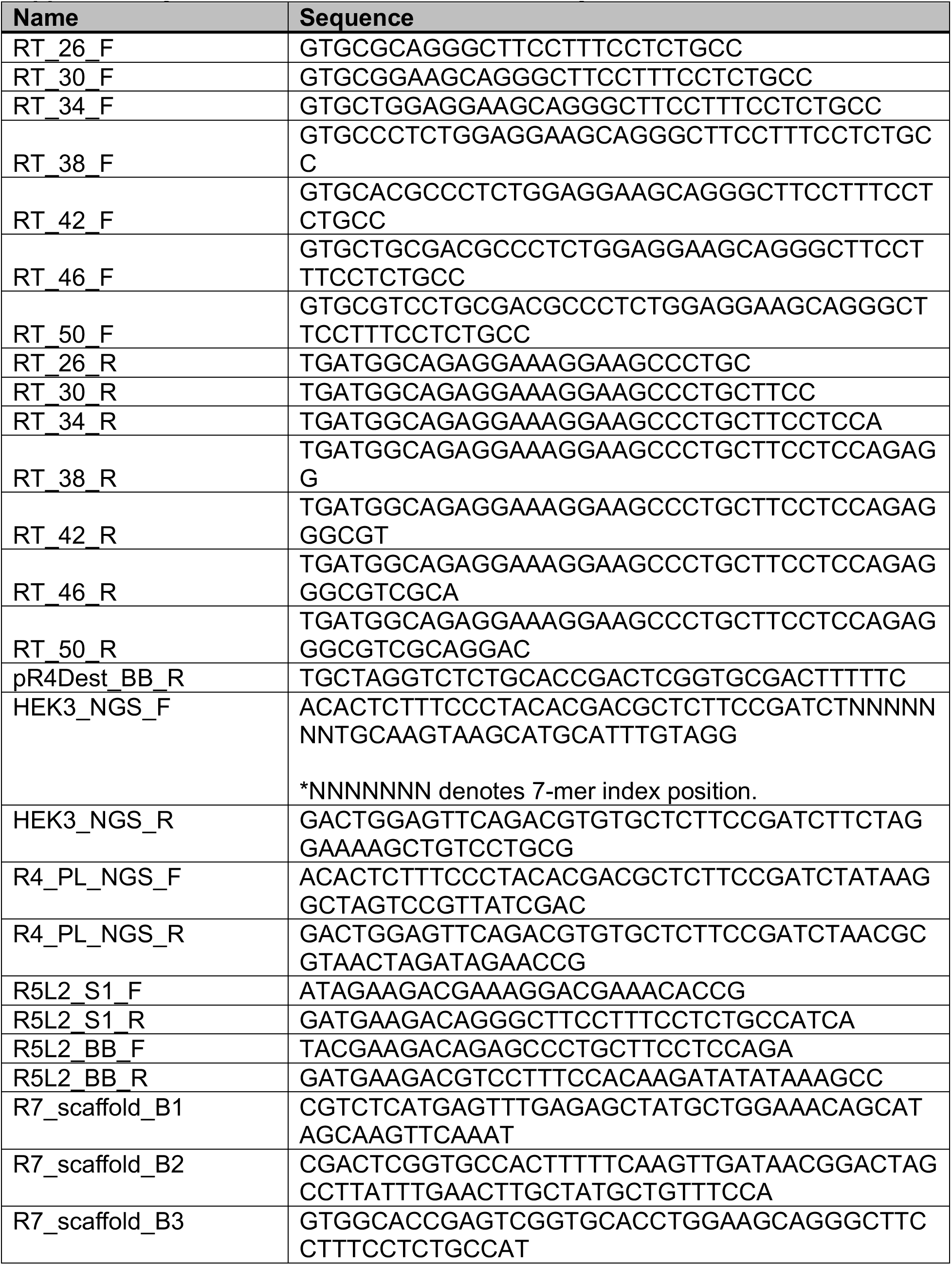
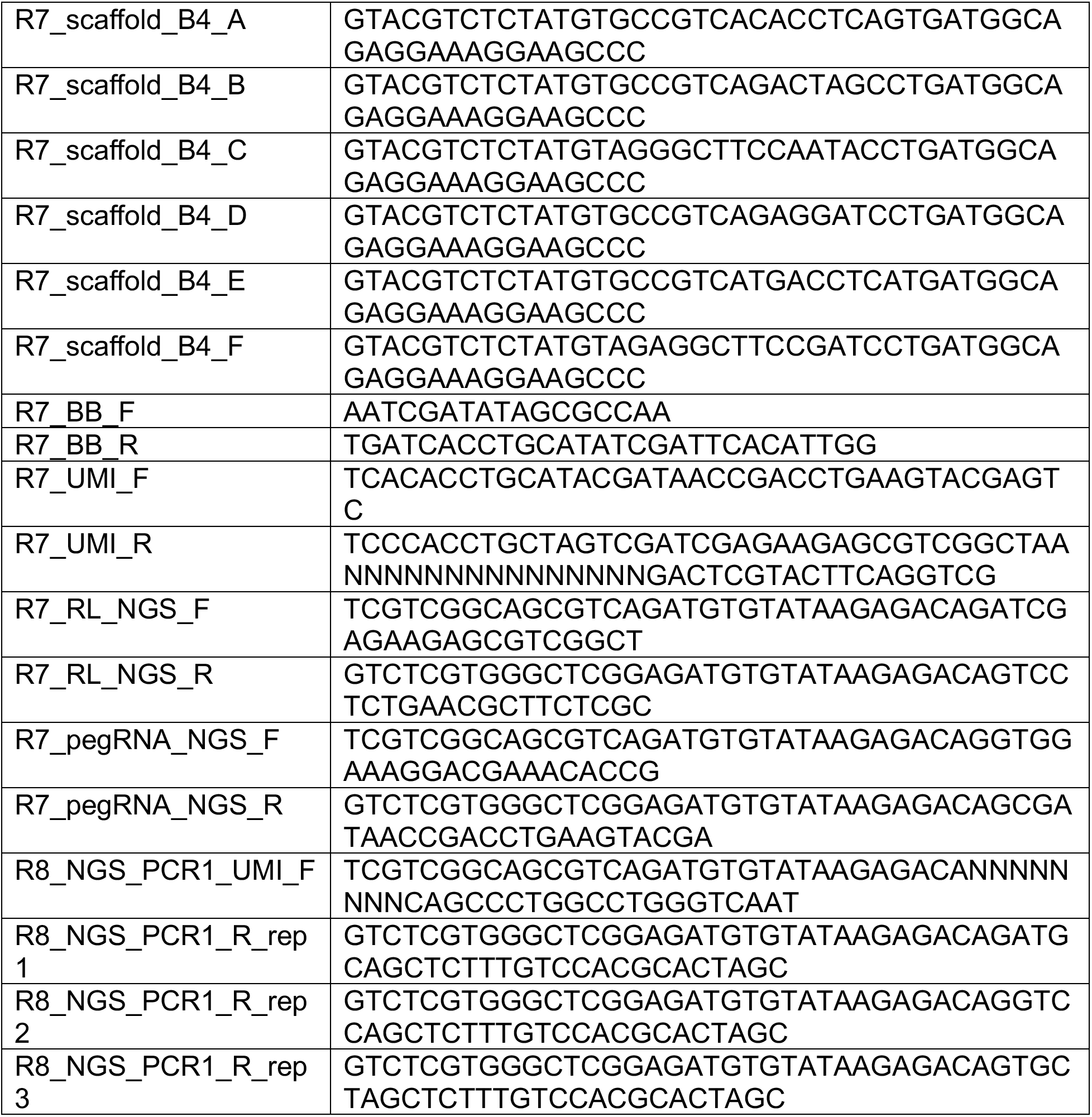
Primers Used in this Study.

**Supplementary Table 2:**
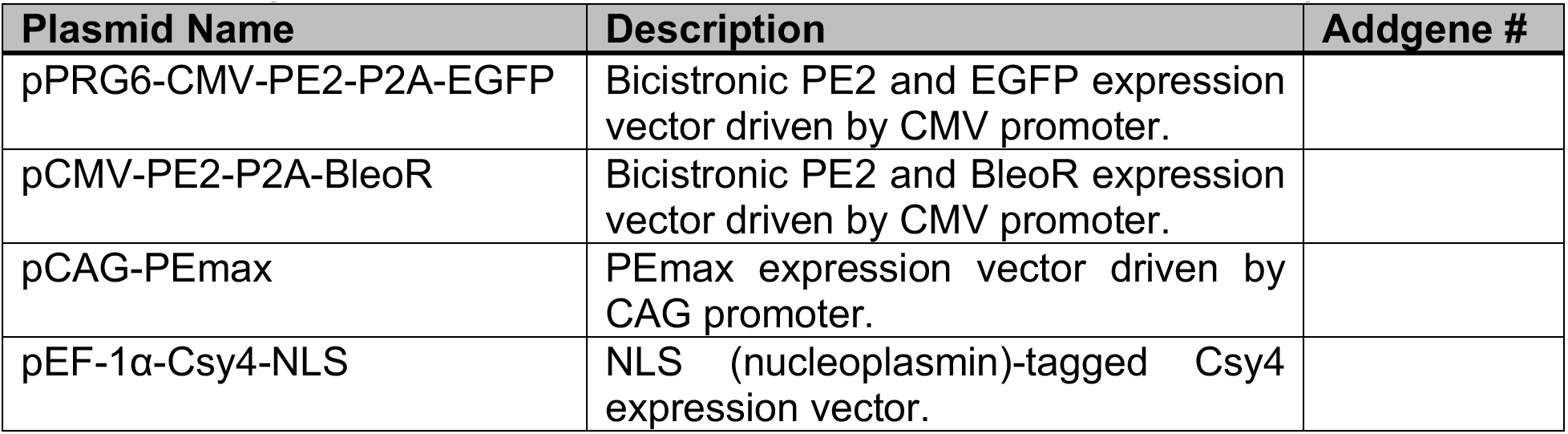
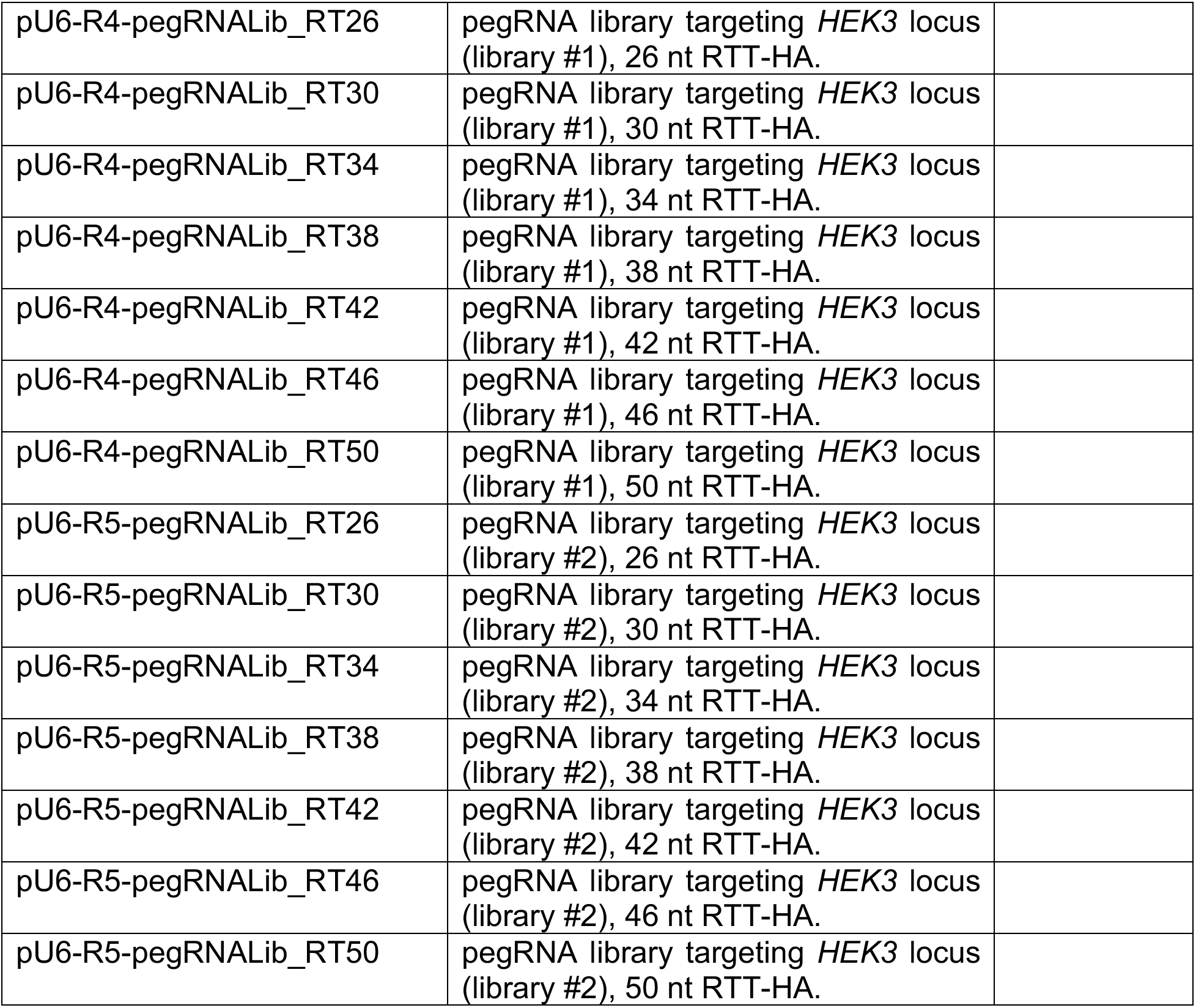
Novel Plasmids / Libraries Used in this Study.

**Supplementary Table 3:**
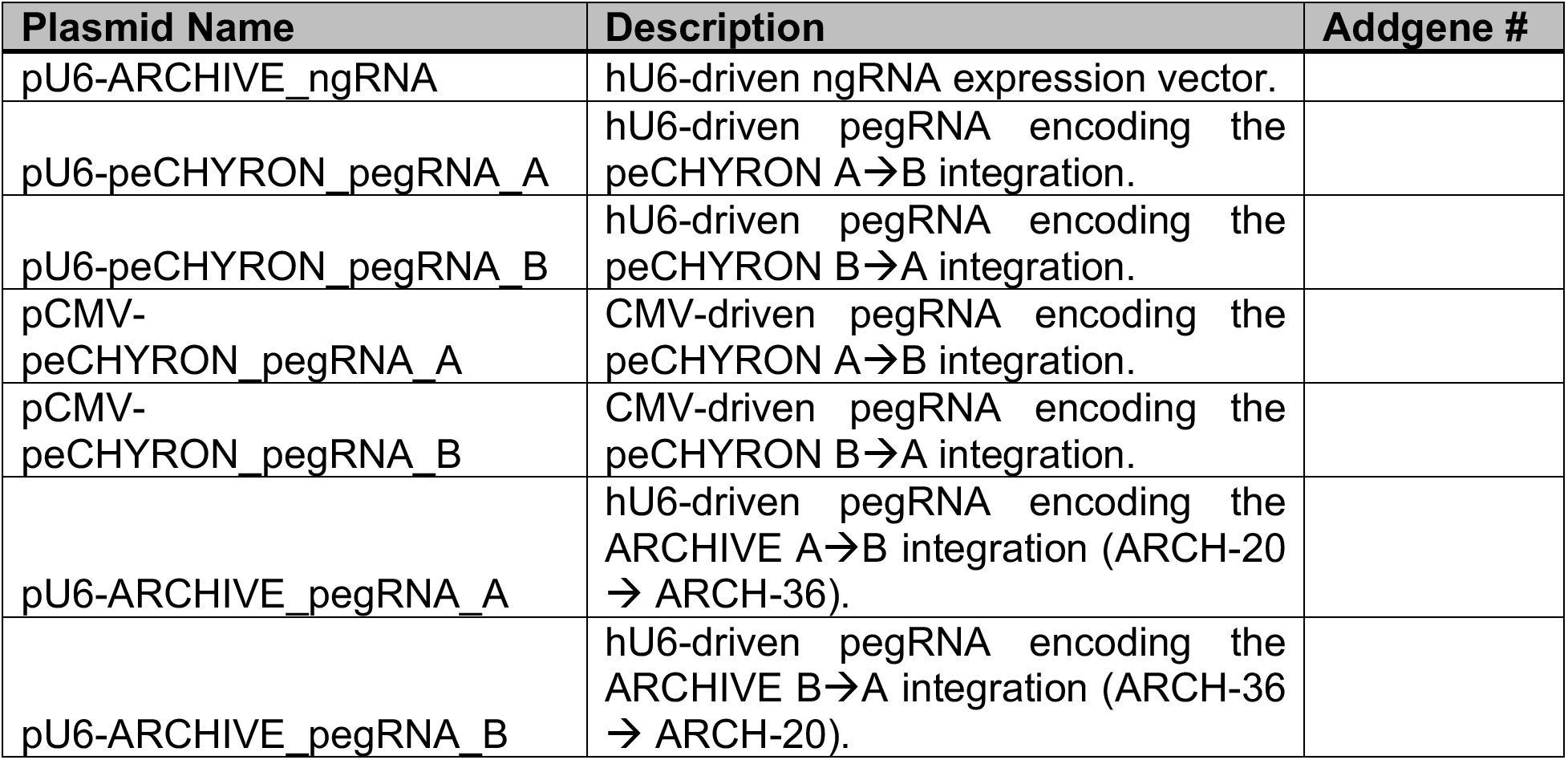
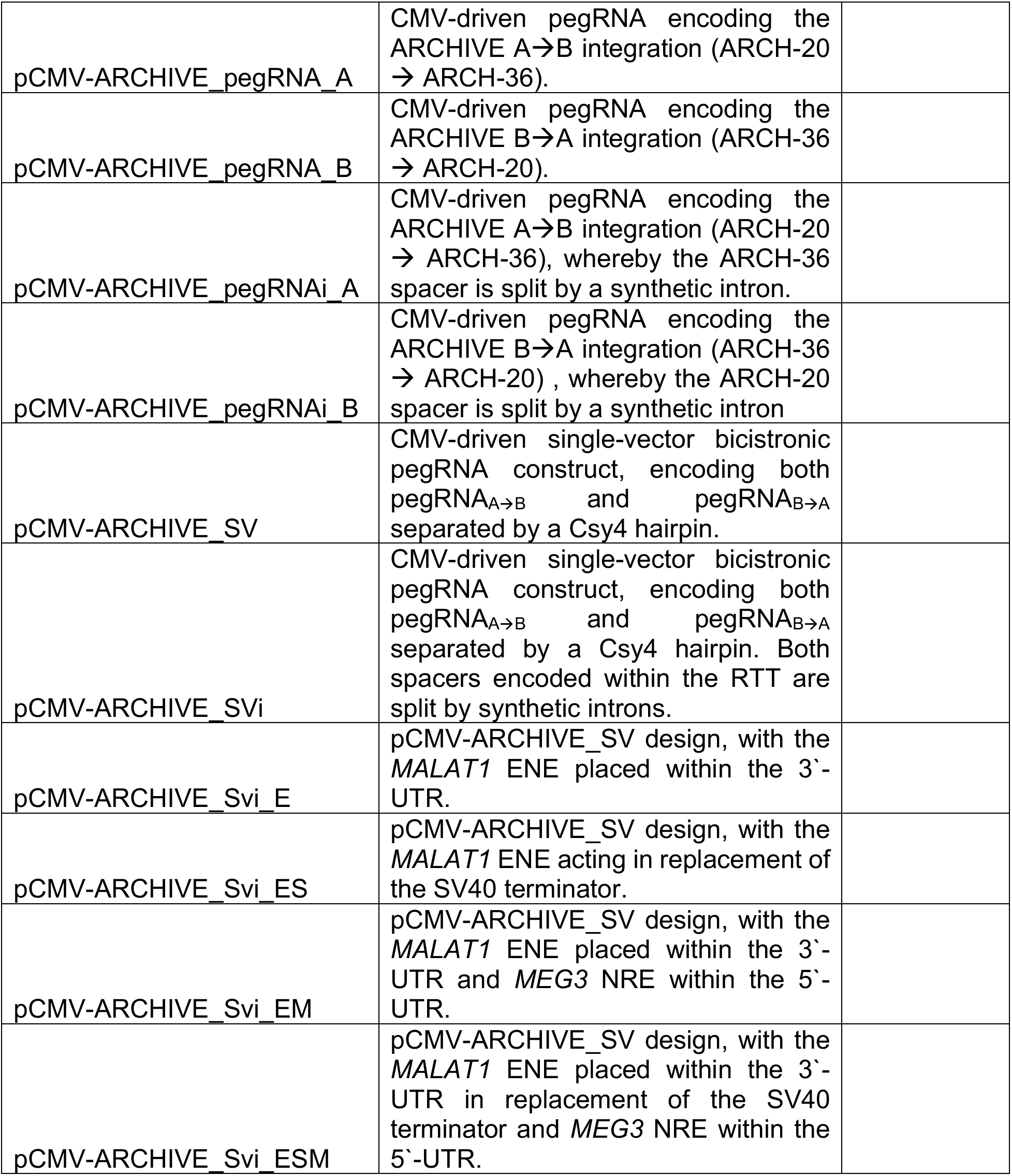
PegRNA Constructs Assembled in this Study.

**Supplementary Table 4:**
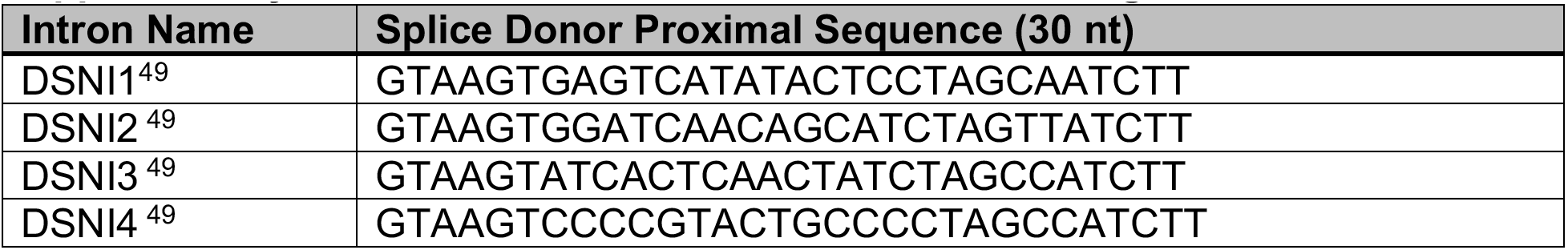
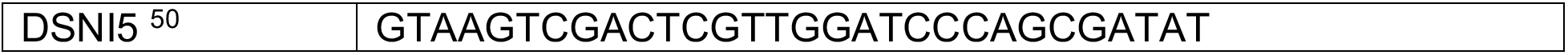
Introns Used for *in Silico* Screening.

## Notes

### Competing Interest Statement

The authors have declared no competing interest.

### Summary of Updates

Edited manuscript body text, results remain unchanged from initial draft.

