## Supplementary Information for "ARCHIVE: An Efficient Open-Ended DNA Recording Device Capable of Multiplex Capture of Pol-II Transcribed Signals"

### Supplementary Data:

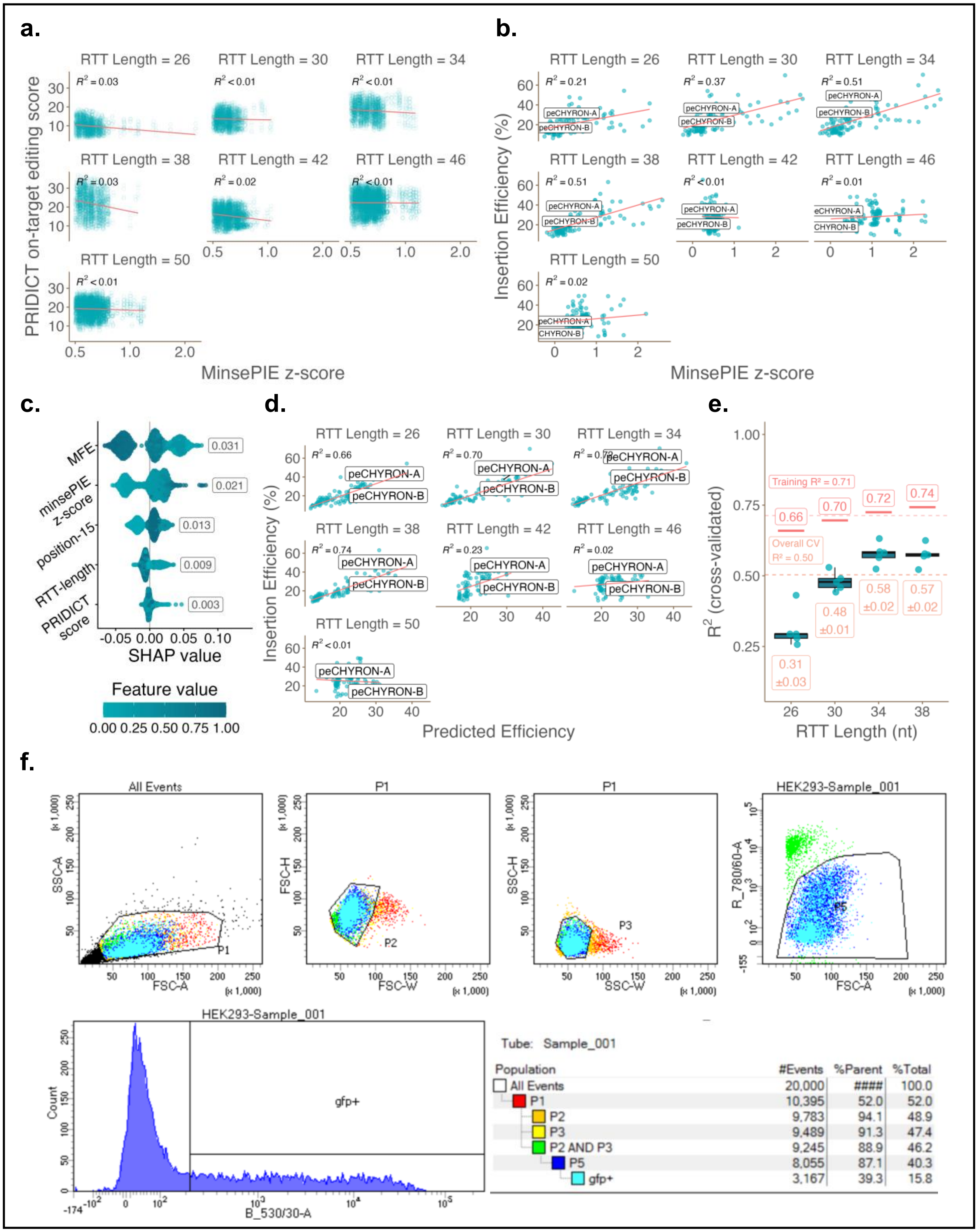

**Supplementary Figure 1: Machine-Guided *in Silico* Evolution of Spacers to Permit High-Efficiency Edit Integration, and Validation Thereof. (a)** Correlation between MinsePIE z-score and PRIDICT on-target score across all evolved spacer sequences, stratified by RTT homology arm length. Line of best fit and $R^2$ are overlayed on each plot facet. **(b)** Correlation between MinsePIE z-score and scaled insertion efficiency across all experimentally validated spacer sequences, stratified by RTT homology arm length. Line of best fit and $R^2$ are overlayed on each plot facet. **(c)** SHAP beeswarm plot of random forest (RF) features (RTT MFE, MinsePIE z-score, spacer position 15 nucleotide, RTT-HA length, and PRIDICT on-target score) used to predict insertion efficiency. Mean absolute SHAP values for each feature are displayed beside each beeswarm respectively. **(d)** Correlation between RF-predicted and experimentally determined insertion efficiency across all tested spacer sequences, stratified by RTT homology arm length. Line of best fit and $R^2$ are overlayed on each plot facet. **(e)** Cross-validated random forest model performance across RTT-HA lengths used for training. Boxplots display the distribution of $R^2$ values across 50 cross-validation splits (5-fold × 10 repeats), stratified by spacer identity to prevent data leakage between RT-length replicates of the same spacer. Red horizontal bars and labels indicate in-sample training $R^2$ per RTT-HA length from the full model; Orange dashed line indicates overall training and cross-validated $R^2$, respectively. Mean cross-validated $R^2$ ± S.E.M. across splits is shown below each group. **(f)** Flow cytometric gating strategy for FACS-sorting HEK293T cells transfected with pegRNA libraries and Subsequent NGS Analysis Workflow. Cells were identified by FSC-A/SSC-A gating, followed by singlet isolation (FSC-W/FSC-H; P2) → (SSC-W/SSC-H; P3) in addition to viability gating (LIVE/DEAD-Near-IR; P5), prior to gating for EGFP+ cells (gfp+).

**Supplementary Table 1: Primers Used in this Study**

| Name | Sequence |
|---|---|
| RT_26_F | GTGCGCAGGGCTTCCTTTCCTCTGCC |
| RT_30_F | GTGCGGAAGCAGGGCTTCCTTTCCTCTGCC |
| RT_34_F | GTGCTGGAGGAAGCAGGGCTTCCTTTCCTCTGCC |
| RT_38_F | GTGCCCTCTGGAGGAAGCAGGGCTTCCTTTCCTCTGC C |
| RT_42_F | GTGCACGCCCTCTGGAGGAAGCAGGGCTTCCTTTCCT CTGCC |
| RT_46_F | GTGCTGCGACGCCCTCTGGAGGAAGCAGGGCTTCCT TTCCTCTGCC |
| RT_50_F | GTGCGTCCTGCGACGCCCTCTGGAGGAAGCAGGGCT TCCTTTCCTCTGCC |
| RT_26_R | TGATGGCAGAGGAAAGGAAGCCCTGC |
| RT_30_R | TGATGGCAGAGGAAAGGAAGCCCTGCTTCC |
| RT_34_R | TGATGGCAGAGGAAAGGAAGCCCTGCTTCCTCCA |
| RT_38_R | TGATGGCAGAGGAAAGGAAGCCCTGCTTCCTCCAGAG G |
| RT_42_R | TGATGGCAGAGGAAAGGAAGCCCTGCTTCCTCCAGAG GGCGT |
| RT_46_R | TGATGGCAGAGGAAAGGAAGCCCTGCTTCCTCCAGAG GGCGTCGCA |
| RT_50_R | TGATGGCAGAGGAAAGGAAGCCCTGCTTCCTCCAGAG GGCGTCGCAGGAC |
| pR4Dest_BB_R | TGCTAGGTCTCTGCACCGACTCGGTGCGACTTTTTC |
| HEK3_NGS_F | ACACTCTTTCCCTACACGACGCTCTTCCGATCTNNNNN NNTGCAAGTAAGCATGCATTTGTAGG<br><br>*NNNNNNN denotes 7-mer index position. |
| HEK3_NGS_R | GACTGGAGTTCAGACGTGTGCTCTTCCGATCTTCTAG GAAAAGCTGTCCTGCG |
| R4_PL_NGS_F | ACACTCTTTCCCTACACGACGCTCTTCCGATCTATAAG GCTAGTCCGTTATCGAC |
| R4_PL_NGS_R | GACTGGAGTTCAGACGTGTGCTCTTCCGATCTAACGC GTAACTAGATAGAACCG |
| R5L2_S1_F | ATAGAAGACGAAAGGACGAAACACCG |
| R5L2_S1_R | GATGAAGACAGGGCTTCCTTTCCTCTGCCATCA |
| R5L2_BB_F | TACGAAGACAGAGCCCTGCTTCCTCCAGA |
| R5L2_BB_R | GATGAAGACGTCCTTTCCACAAGATATATAAAGCC |
| R7_scaffold_B1 | CGTCTCATGAGTTTGAGAGCTATGCTGGAAACAGCAT AGCAAGTTCAAAT |
| R7_scaffold_B2 | CGACTCGGTGCCACTTTTTCAAGTTGATAACGGACTAG CCTTATTTGAACTTGCTATGCTGTTTCCA |
| R7_scaffold_B3 | GTGGCACCGAGTCGGTGCACCTGGAAGCAGGGCTTC CTTTCCTCTGCCAT |

| | |
|---|---|
| R7_scaffold_B4_A | GTACGTCTCTATGTGCCGTCACACCTCAGTGATGGCAGAGGAAAGGAAGCCC |
| R7_scaffold_B4_B | GTACGTCTCTATGTGCCGTCAGACTAGCCTGATGGCAGAGGAAAGGAAGCCC |
| R7_scaffold_B4_C | GTACGTCTCTATGTAGGGCTTCCAATACCTGATGGCAGAGGAAAGGAAGCCC |
| R7_scaffold_B4_D | GTACGTCTCTATGTGCCGTCAGAGGATCCTGATGGCAGAGGAAAGGAAGCCC |
| R7_scaffold_B4_E | GTACGTCTCTATGTGCCGTCATGACCTCATGATGGCAGAGGAAAGGAAGCCC |
| R7_scaffold_B4_F | GTACGTCTCTATGTAGAGGCTTCCGATCCTGATGGCAGAGGAAAGGAAGCCC |
| R7_BB_F | AATCGATATAGCGCCAA |
| R7_BB_R | TGATCACCTGCATATCGATTCACATTGG |
| R7_UMI_F | TCACACCTGCATACGATAACCGACCTGAAGTACGAGTC |
| R7_UMI_R | TCCCACCTGCTAGTCGATCGAGAAGAGCGTCGGCTAANNNNNNNNNNNNNNNGACTCGTACTTCAGGTCG |
| R7_RL_NGS_F | TCGTCGGCAGCGTCAGATGTGTATAAGAGACAGATCGAGAAGAGCGTCGGCT |
| R7_RL_NGS_R | GTCTCGTGGGCTCGGAGATGTGTATAAGAGACAGTCCTCTGAACGCTTCTCGC |
| R7_pegRNA_NGS_F | TCGTCGGCAGCGTCAGATGTGTATAAGAGACAGGTGGAAAGGACGAAACACCG |
| R7_pegRNA_NGS_R | GTCTCGTGGGCTCGGAGATGTGTATAAGAGACAGCGATAACCGACCTGAAGTACGA |
| R8_NGS_PCR1_UMI_F | TCGTCGGCAGCGTCAGATGTGTATAAGAGACANNNNNNNNCAGCCCTGGCCTGGGTCAAT |
| R8_NGS_PCR1_R_rep1 | GTCTCGTGGGCTCGGAGATGTGTATAAGAGACAGATGCAGCTCTTTGTCCACGCACTAGC |
| R8_NGS_PCR1_R_rep2 | GTCTCGTGGGCTCGGAGATGTGTATAAGAGACAGGTCCAGCTCTTTGTCCACGCACTAGC |
| R8_NGS_PCR1_R_rep3 | GTCTCGTGGGCTCGGAGATGTGTATAAGAGACAGTGCTAGCTCTTTGTCCACGCACTAGC |

**Supplementary Table 2: Novel Plasmids / Libraries Used in this Study**

| Plasmid Name | Description | Addgene # |
|---|---|---|
| pPRG6-CMV-PE2-P2A-EGFP | Bicistronic PE2 and EGFP expression vector driven by CMV promoter. | |
| pCMV-PE2-P2A-BleoR | Bicistronic PE2 and BleoR expression vector driven by CMV promoter. | |
| pCAG-PEmax | PEmax expression vector driven by CAG promoter. | |
| pEF-1α-Csy4-NLS | NLS (nucleoplasmin)-tagged Csy4 expression vector. | |

| | |
|---|---|
| pU6-R4-pegRNALib_RT26 | pegRNA library targeting *HEK3* locus (library #1), 26 nt RTT-HA. |
| pU6-R4-pegRNALib_RT30 | pegRNA library targeting *HEK3* locus (library #1), 30 nt RTT-HA. |
| pU6-R4-pegRNALib_RT34 | pegRNA library targeting *HEK3* locus (library #1), 34 nt RTT-HA. |
| pU6-R4-pegRNALib_RT38 | pegRNA library targeting *HEK3* locus (library #1), 38 nt RTT-HA. |
| pU6-R4-pegRNALib_RT42 | pegRNA library targeting *HEK3* locus (library #1), 42 nt RTT-HA. |
| pU6-R4-pegRNALib_RT46 | pegRNA library targeting *HEK3* locus (library #1), 46 nt RTT-HA. |
| pU6-R4-pegRNALib_RT50 | pegRNA library targeting *HEK3* locus (library #1), 50 nt RTT-HA. |
| pU6-R5-pegRNALib_RT26 | pegRNA library targeting *HEK3* locus (library #2), 26 nt RTT-HA. |
| pU6-R5-pegRNALib_RT30 | pegRNA library targeting *HEK3* locus (library #2), 30 nt RTT-HA. |
| pU6-R5-pegRNALib_RT34 | pegRNA library targeting *HEK3* locus (library #2), 34 nt RTT-HA. |
| pU6-R5-pegRNALib_RT38 | pegRNA library targeting *HEK3* locus (library #2), 38 nt RTT-HA. |
| pU6-R5-pegRNALib_RT42 | pegRNA library targeting *HEK3* locus (library #2), 42 nt RTT-HA. |
| pU6-R5-pegRNALib_RT46 | pegRNA library targeting *HEK3* locus (library #2), 46 nt RTT-HA. |
| pU6-R5-pegRNALib_RT50 | pegRNA library targeting *HEK3* locus (library #2), 50 nt RTT-HA. |

**Supplementary Table 3: PegRNA Constructs Assembled in this Study**

| Plasmid Name | Description | Addgene # |
|---|---|---|
| pU6-ARCHIVE_ngRNA | hU6-driven ngRNA expression vector. | |
| pU6-peCHYRON_pegRNA_A | hU6-driven pegRNA encoding the peCHYRON A→B integration. | |
| pU6-peCHYRON_pegRNA_B | hU6-driven pegRNA encoding the peCHYRON B→A integration. | |
| pCMV-peCHYRON_pegRNA_A | CMV-driven pegRNA encoding the peCHYRON A→B integration. | |
| pCMV-peCHYRON_pegRNA_B | CMV-driven pegRNA encoding the peCHYRON B→A integration. | |
| pU6-ARCHIVE_pegRNA_A | hU6-driven pegRNA encoding the ARCHIVE A→B integration (ARCH-20 → ARCH-36). | |
| pU6-ARCHIVE_pegRNA_B | hU6-driven pegRNA encoding the ARCHIVE B→A integration (ARCH-36 → ARCH-20). | |

| | |
|---|---|
| pCMV-ARCHIVE_pegRNA_A | CMV-driven pegRNA encoding the ARCHIVE A→B integration (ARCH-20 → ARCH-36). |
| pCMV-ARCHIVE_pegRNA_B | CMV-driven pegRNA encoding the ARCHIVE B→A integration (ARCH-36 → ARCH-20). |
| pCMV-ARCHIVE_pegRNAi_A | CMV-driven pegRNA encoding the ARCHIVE A→B integration (ARCH-20 → ARCH-36), whereby the ARCH-36 spacer is split by a synthetic intron. |
| pCMV-ARCHIVE_pegRNAi_B | CMV-driven pegRNA encoding the ARCHIVE B→A integration (ARCH-36 → ARCH-20) , whereby the ARCH-20 spacer is split by a synthetic intron |
| pCMV-ARCHIVE_SV | CMV-driven single-vector bicistronic pegRNA construct, encoding both pegRNA$_{A \rightarrow B}$ and pegRNA$_{B \rightarrow A}$ separated by a Csy4 hairpin. |
| pCMV-ARCHIVE_SVi | CMV-driven single-vector bicistronic pegRNA construct, encoding both pegRNA$_{A \rightarrow B}$ and pegRNA$_{B \rightarrow A}$ separated by a Csy4 hairpin. Both spacers encoded within the RTT are split by synthetic introns. |
| pCMV-ARCHIVE_Svi_E | pCMV-ARCHIVE_SV design, with the *MALAT1* ENE placed within the 3`-UTR. |
| pCMV-ARCHIVE_Svi_ES | pCMV-ARCHIVE_SV design, with the *MALAT1* ENE acting in replacement of the SV40 terminator. |
| pCMV-ARCHIVE_Svi_EM | pCMV-ARCHIVE_SV design, with the *MALAT1* ENE placed within the 3`-UTR and *MEG3* NRE within the 5`-UTR. |
| pCMV-ARCHIVE_Svi_ESM | pCMV-ARCHIVE_SV design, with the *MALAT1* ENE placed within the 3`-UTR in replacement of the SV40 terminator and *MEG3* NRE within the 5`-UTR. |

**Supplementary Table 4: Introns Used for *in Silico* Screening**

| Intron Name | Splice Donor Proximal Sequence (30 nt) |
|---|---|
| DSNI1[44] | GTAAGTGAGTCATATACTCCTAGCAATCTT |
| DSNI2[44] | GTAAGTGGATCAACAGCATCTAGTTATCTT |
| DSNI3[44] | GTAAGTATCACTCAACTATCTAGCCATCTT |
| DSNI4[44] | GTAAGTCCCCGTACTGCCCCTAGCCATCTT |

| | |
|---|---|
| DSNI5 [45] | GTAAGTCGACTCGTTGGATCCCAGCGATAT |
